# 3D-intrusions transport active surface microbial assemblages to the dark ocean

**DOI:** 10.1101/2023.09.14.557835

**Authors:** Mara A. Freilich, Camille Poirier, Mathieu Dever, Eva Alou-Font, John Allen, Andrea Cabornero, Lisa Sudek, Chang Jae Choi, Simón Ruiz, Ananda Pascual, J. Thomas Farrar, T.M. Shaun Johnston, Eric D’Asaro, Alexandra Z. Worden, Amala Mahadevan

## Abstract

Subtropical oceans contribute significantly to global primary production, but the fate of the picophytoplankton that dominate in these low nutrient regions is poorly understood. Working in the subtropical Mediterranean, we demonstrate that subduction of water at ocean fronts generates 3D intrusions with uncharacteristically high carbon, chlorophyll, and oxygen that extend below the sunlit photic-zone into the dark ocean. These contain “fresh” picophytoplankton assemblages that resemble the photic-zone regions where the water originated. Intrusions propagate depth-dependent seasonal variations in microbial assemblages into the ocean interior. Strikingly, the intrusions included dominant biomass contributions from non-photosynthetic bacteria and enrichment of enigmatic heterotrophic bacterial lineages. Thus, the intrusions not only deliver material that differs in composition and nutritional character from sinking detrital particles, but also drive shifts in bacterial community composition, organic matter processing, and interactions between surface and deep communities. Modeling efforts paired with global observations demonstrate that subduction can flux similar magnitudes of particulate organic carbon as sinking export, but is not accounted for in current export estimates and carbon cycle models. Intrusions formed by subduction are a particularly important mechanism for enhancing connectivity between surface and upper mesopelagic ecosystems in stratified subtropical ocean environments that are expanding due to the warming climate.

## Introduction

Surface ocean carbon fixation by photosynthetic planktonic microbes generates particulate organic carbon (POC) that fuels marine ecosystems^1^. A portion of this POC is exported to the underlying dark ocean where it is a critical energy source for heterotrophic organisms. However, particularly in the subtropics, the mechanisms for export of POC are unclear^2,3^ despite the fact that a major fraction of global primary production occurs in these regions^1^. In subtropical regions, the sunlit photic zone is dominated by picophytoplankton (diameters <3µm) that are non-sinking due to their small sizes. These unicellular microbes include the picocyanobacteria, *Prochlorococcus* and *Synechococcus*, as well as picoeukaryotes, and succeed in oligotrophic subtropical gyres because their small size is advantageous for nutrient acquisition under limiting conditions^4,5^. Importantly, picophytoplankton-dominated environments are expanding due to surface ocean warming, which strengthens vertical stratification and induces nutrient-limiting conditions^6,7^.

Contrary to the expectation that picophytoplankton cells would not arrive intact in the dark mesopelagic (depths ≥200m) due to the hundreds of days it would take for them to sink, DNA, photosynthetic pigments (e.g., chlorophyll *a*), or cells of cyanobacteria have occasionally been found in subtropical mesopelagic sediment traps and samples^8–10^. Various mechanisms for picophytoplankton packaging, including proposed aggregation of dead cells during viral lysis or in zooplankton fecal matter are among explanations for enhanced export efficiency. These mechanisms may explain some of the observed picophytoplanktonic material at dark depths, depending on degradation rates and other factors. Additionally, physical processes involving movement of water masses, such as polar thermohaline-driven sinking^11^, and advection by oceanic surface mixed layer eddies^12^, result in export of picophytoplankton and oxygen from the surface. The surface-enhanced submesoscale (1-10 km scales) dynamics that generate the mixed layer eddies and filaments needed for the latter are found in regions with deep surface mixed layers and strong surface forcing, such as the subpolar North Atlantic, coastal upwelling fronts, and the Southern Ocean^12,13^. However, the shallowness of the mixed layer in the subtropics results in weaker submesoscale dynamics^14^. Further, since subtropical surface waters are nutrient-depleted, phytoplankton biomass is concentrated below the mixed layer, making it less prone to export by mixed layer processes.

By contrast, mesoscale processes (10-100 km scale) that generate sloping density surfaces (fronts) are ubiquitous in the subtropics and can facilitate relatively rapid submesoscale vertical motion (order 100 m day^-1^) but their contributions to POC export from the photic zone are generally not considered. Building on prior understanding of the impact of frontal dynamics on primary production, mesoscale vertical motion, and vertical transport of chlorophyll in the subtropical Mediterranean Sea^16–18^, we undertook advanced physical oceanographic measurements to identify signatures of active submesoscale vertical exchanges in real-time and analyzed resident microbial communities using flow cytometry and molecular analyses. Together, these observations enabled us to trace 3-dimensional chlorophyll-containing intrusions to the upper mesopelagic and determine their environmental origins in the photic zone (upper 70 m). We observed intact picophytoplankton cells in the aphotic zone at mesoscale fronts in this region. We then probed differentiated microbial assemblages and quantified their contributions to carbon export in the upper mesopelagic. The results show that alongside photosynthetic cells, a surprisingly large proportion of heterotrophic bacteria, and their carbon, are transported to the dark ocean via the 3D-intrusions in a non-decayed state. Moreover, two enigmatic, broadly distributed marine bacterial groups, Marinimicrobia (SAR406) and Parvibaculales (OCS116)^19–21^, appear to be stimulated in the intrusions.

Drawing together these lines of evidence, we demonstrate that intrusions enhance picoplankton carbon export and alter microbial diversity and composition through surface-to-mesopelagic pathways. The varied outcomes of microbial interactions in the upper mesopelagic may allow for enriched genetic and metabolite exchanges between microbial taxa from different habitats, alongside delivering POC that differs in nutritional composition and lability from ‘background’ mesopelagic waters. Upon establishing the importance of this mechanism for picoplankton export and community structure, we explored its potential significance at a global level, finding that intrusion dynamics contribute as much carbon to the upper mesopelagic as estimated sinking in subtropical regions.

### Chlorophyll-containing intrusions in the dark ocean are formed by frontal subduction

While subduction of chlorophyll to the upper mesopelagic has been documented in the Western Mediterranean Sea^16,18^, the biophysical dynamics and ecological significance of these chlorophyll features are not understood. Therefore, we conducted ship-based expeditions in July 2017, May-June 2018, and March-April 2019 wherein oceanic density fronts in the Western Mediterranean Sea were repeatedly transected (Fig. 1a). Fronts in this region partition relatively fresh, cool and buoyant Atlantic waters that enter the Alborán Sea, from saltier, warmer, and denser Mediterranean waters (Extended Data Fig. 1). We measured salinity, temperature, and bio-optical properties at approximately every meter from the surface to 200 m depth, using towed profilers. This was performed as the ship transited orthogonal to strong frontal flows that separate the Western and Eastern Alborán gyres and are known to evolve on timescales of weeks to months^22^.

**Fig. 1.**
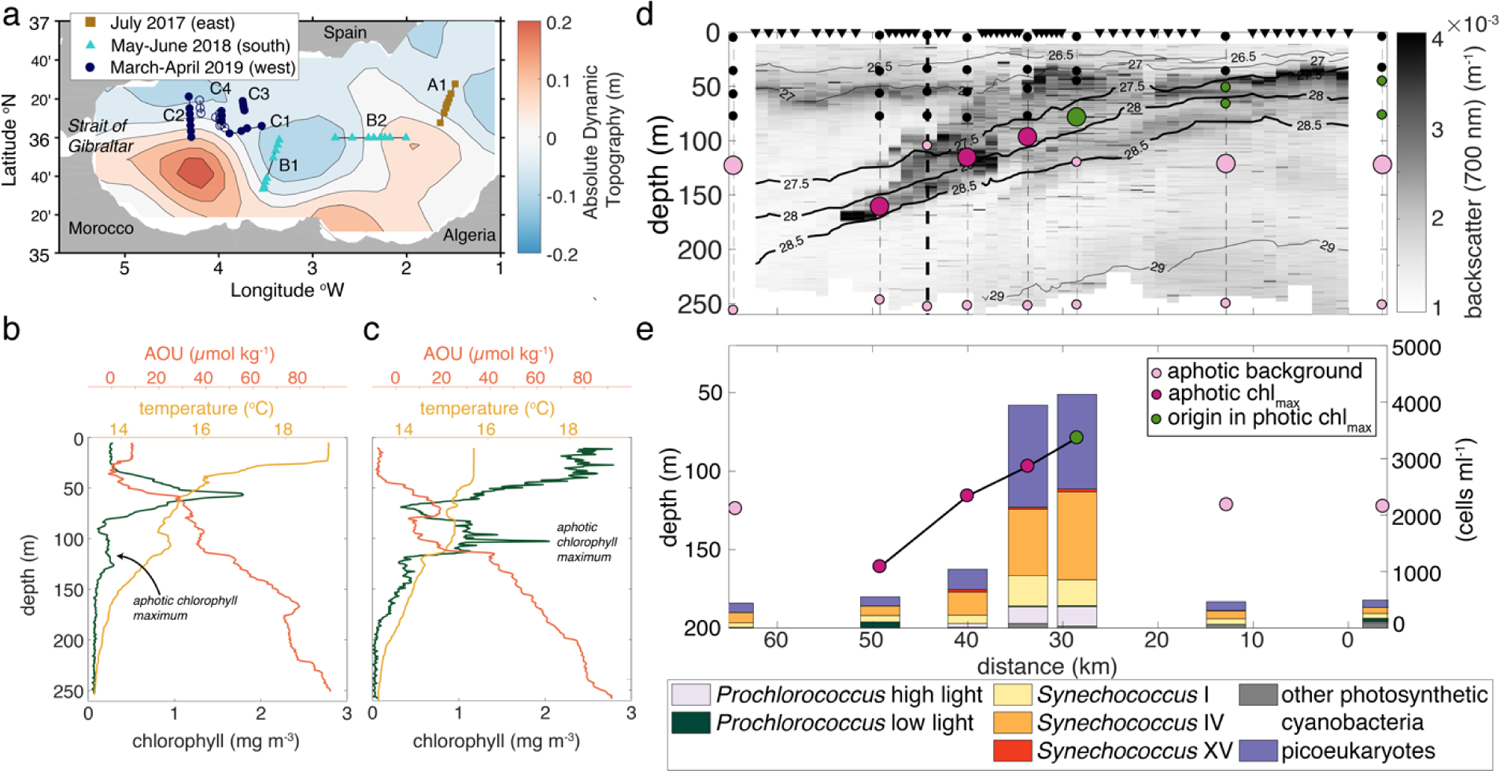
Chlorophyll maxima below the Mediterranean Sea photic zone are formed by surface-derived intrusions. **a,** Eastern and Western Alborán gyres and transects (lines) sampled synoptically (symbols) and locations where molecular analyses and flow cytometry were performed (filled symbols); Background shows absolute dynamic topography (ADT) on May 25, 2018, illustrating general gyre positioning. **b,c,** Chlorophyll fluorescence, apparent oxygen utilization (AOU), and temperature for representatives of the two profile types in summer (May 2018) **(b)** and winter/spring (March 2019) **(c)**, with the aphotic chlorophyll maxima (indicated) between 100-150 m and co-occurring inversions relative to background trends in temperature and AOU. **d,** Example 2-dimensional section across transect B2 (panel B), with 0 km being at the western edge. Locations of EcoCTD casts (triangles), standard CTD casts (dashed lines) and backscatter (grey scale) indicated. The panel C cast is highlighted for reference (bold dashed line). Molecular samples (dots) originating from the photic zone (black dots), ACM (dark pink), origin waters of the ACM, namely the chlorophyll maximum on the dense side of the front (green), or the aphotic background (light pink); larger dots represent those displayed in panel F. **e,** Phytoplankton abundances and community composition by V1-V2 16S rRNA gene ASV analyses along transect B2 at depths representing the intrusion (green and dark pink connected by a line) and in background samples (light pink; i.e. same depth as intrusions but outside).

Our measurements demonstrated that in the regions and years sampled the chlorophyll and biomass maxima coincide within the photic zone (Fig. 1b, c, Extended Data Figs. 2, 3), based on *in vivo* chlorophyll *a* fluorescence and optical backscatter at 700 nm (a proxy for POC), respectively, as observed previously^23^. This established that peaks in chlorophyll fluorescence resulted from increased biomass, as opposed to phytoplankton photoacclimation. More generally, two types of ‘classical’ water column structures were observed within the photic zone: (i) a deep chlorophyll maximum (DCM) at the top of the nutricline and beneath the mixed layer (Fig. 1b; mixed layer depth 5–20 m and 1% light level 50–60 m) during warmer periods when the photic zone exhibits thermal stratification, as observed in May-June 2017 and July 2018; and (ii) a well-mixed surface layer with relatively even chlorophyll distributions between 0 and 70 m during the March-April 2019 expedition (Fig. 1c; 1% light level was 60–70m). Such seasonal patterns in the vertical structure are similar to those of the Atlantic and Pacific subtropical gyres^24^.

Strikingly, on multiple transects, we also observed secondary maxima in chlorophyll within the aphotic zone (deeper than 90 m), beneath the seasonally varying surface mixed layer as identified by uniform density (e.g., Fig. 1b, 1c). The secondary, aphotic chlorophyll maxima (ACM) co-occurred with secondary maxima in optical backscatter at 700 nm and beam transmission (another proxy for POC), alongside secondary minima in apparent oxygen utilization (AOU) or, equivalently, maxima in oxygen saturation, and temperature-salinity (spice) anomalies (Fig. 1b-d). The concurrent ACM and minima in AOU were indicative of water being subducted from the photic zone on time scales short enough that oxygen levels remained high and fluorescing chlorophyll was present in the dark.

The anomalous chlorophyll features and associated water mass characteristics were inconsistent with mechanisms that have been previously suggested for ACM formation in other subtropical regions. Photoautotrophic growth is not sufficient to drive the observations, given the lack of sunlight, and because the co-occurring peaks in chlorophyll and POC indicate that phytoplankton photoacclimation did not generate the maxima (Extended Data Fig. 4). Likewise, vertical mixing does not generate secondary maxima or minima or the observed temperature-salinity anomalies. Moreover, sinking POC would not generate the anomalously high oxygen concentrations observed below the photic zone. Thus, the ACMs, which are seen in contiguous profiles at the same (or similar) density appear to be along-isopycnal intrusions.

To pursue the biological dynamics and origins of the observed ACMs, we examined the nature of the material contributing to ACM signatures in a biological and ecological framework. To this end, we turned to the full complement of biological, physical, and bio-optical measurements recovered at a horizontal spacing of 1–10 km on 10 frontal crossings (Fig. 1a), which offers a highly resolved picture of microbial community variability. By observing downward CTD cast data in real-time, we adaptively sampled 6 depths during the upcast, which included (a) the ACM, when present, (b) regions of high chlorophyll *a* and biomass in the photic zone; either the well-mixed high chlorophyll layer within the surface mixed layer or DCM in the stratified region below the surface mixed layer, (c) photic zone depths not within a chlorophyll maximum, and (d) aphotic, upper mesopelagic background samples from below the photic zone (but not within an ACM). Our sampling also targeted consistent density surfaces on a given cross-frontal transect, when possible, as advective transport occurs along isopycnals, or surfaces of constant density. Cell enumeration by flow cytometry in these features consistently demonstrated that both photosynthetic eukaryotes and cyanobacteria had higher abundances in the ACM compared to the ‘background’ upper mesopelagic water, as exemplified in Transect B2 (Fig. 1d,e, Extended Data 5). The combined biological and biophysical analyses of cross frontal transects, showing co-occurrence of ACMs and AOU minima in contiguous vertical profiles, alongside corresponding changes in phytoplankton abundances, suggest they are part of coherent intrusions that provide a 3D pathway for transport from the photic zone to the dark ocean.

The anomalies that characterized intrusions became weaker with increasing depth, specifically of chlorophyll and AOU, as did the temperature-salinity (spice) anomalies, although to a lesser degree (e.g., Fig. 1d,e, Extended Data Fig. 5). Notably, the proportion of cyanobacteria amongst the picophytoplankton increased with depth relative to picoeukaryotes (t-test p = 0.0023, confidence interval = [1.0623, 1.2598] where 1 is no change). This may be attributable to a differential loss of picoeukaryotes due to higher mortality rates^5^, or possibly greater persistence of picocyanobacteria in dark conditions^25^. Changes in biological properties with depth within intrusions relates to the decay of chlorophyll, and respiration of organic matter, as well as mixing with surrounding waters. In contrast, salinity and temperature, which contribute to spice, are preserved and only affected by mixing. Despite weakening signatures, we found that phytoplankton within the intrusions comprise intact cells with classical fluorescence and light scattering properties that differ from dead picophytoplankton cells and aphotic communities outside intrusions. Thus, while modifications in intrusion signatures indicate that deeper parts of intrusions are older, having left the photic zone prior to shallower parts, we also infer from the intact cells that there is a relatively rapid transport in the intrusions, which connect photic zone communities to the mesopelagic on timescales of days.

Our findings point to phytoplankton export on spatial and temporal scales that are unlikely to be achieved through mesoscale-driven vertical transport alone, which was previously used to explain chlorophyll intrusions in the Mediterranean Sea. Intrusions are 3D features that appeared to originate in 1–10 km patches along the fronts and extend downward at least 200m in the vertical and 10s of kilometers transversely, along the sloping density surfaces that constitute the front. The features can extend approximately 100 km in the direction of the front because the along-front velocity is about 1000 times larger than the vertical velocity. This spatial structure is emblematic of the coupling between surface-enhanced submesoscale processes and deeper mesoscale dynamics. This coupling can locally intensify fronts and generate more rapid subduction, as established by our modeling efforts herein (Extended Data Text 9) and previously^18,26^. Together, the modeling studies demonstrate downward instantaneous vertical velocities of up to 100m day^-1^, even below the surface, generating sustained time-integrated vertical transport in intrusions in excess of ∼10m day^-1^ ^18,26,27^. The mechanism identified in our subtropical study region therefore results in transport to greater depths than the wintertime mixed layer in both winter and summer. This prevents remixing to the surface at the annual onset of deep mixing, leading to year-round episodic export of picophytoplankton to the upper mesopelagic. Collectively, the intrusions characterized herein are generated by different physical processes than those previously invoked for small phytoplankton export, which rely on mixed layer processes and transport POC only as deep as the wintertime mixed layer depth^12^.

### Intrusions transport picophytoplankton communities from surface or DCM waters to the upper mesopelagic

We next aimed to more precisely identify the origins of the picophytoplankton communities that reached aphotic depths within the intrusions. To this end, we examined phytoplankton community composition in 221 transect samples using phylogenetic analysis of V1-V2 16S rRNA gene Amplicon Sequence Variants (ASVs), which capture cyanobacteria and plastids from eukaryotic phytoplankton^24,28^, alongside phytoplankton cell abundances. As expected, picophytoplankton groups in the photic zone exhibited both depth-related patterns and seasonality in cellular abundances and community composition related to the seasonal changes in water column structure on our transects and consistent with patterns reported in the Western Mediterranean Sea and other subtropical locations^24,28–30^. With respect to intrusions, water column structure appeared to influence the depth of origin. For example, in Transect B2, which came from a stratified period (May-June, Fig. 1d), communities from the DCM, rather than the surface, appeared to be the origins of the ACM picophytoplankton cells/POC. Specifically, *Synechococcus* clades I and IV and eukaryotic cells were the dominant picophytoplankton in the ACM and their relative abundances were similar in the DCM and ACM (Fig. 1e), while the surface community was dominated by *Synechococcus* clade IV. In contrast, in March-April 2019, when the mixed layer was near its annual maximum depth, the intrusion that generated the ACM appears to be sourced from the surface mixed layer, with the ACM and surface communities being similar (Extended Data Fig. 5). Additionally, high-light adapted ecotypes of *Prochlorococcus*^4^ were present in the ACM at ∼100m, corroborating our finding that the intrusion waters originated in the top layers of the photic zone.

Across all sampled transects, intrusions were more similar to surface communities than background waters, despite the decay in the ‘strength’ of intrusion signals during descent (Fig. 1e, Extended Data Fig. 5), which potentially complicates identification of source waters. Hierarchical clustering based on community composition grouped communities sampled from the ACM primarily with microbial communities from the well-mixed surface layer or DCM (when present) on the dense side of fronts (Fig. 2a). This suggests that intrusions connect chlorophyll maxima in the photic zone to the dark ocean, regardless of the particular depth of that maximum. Due to differences in origin communities from transect to transect, the totality of intrusion communities does not collectively converge to a single unique community composition.

**Fig. 2.**
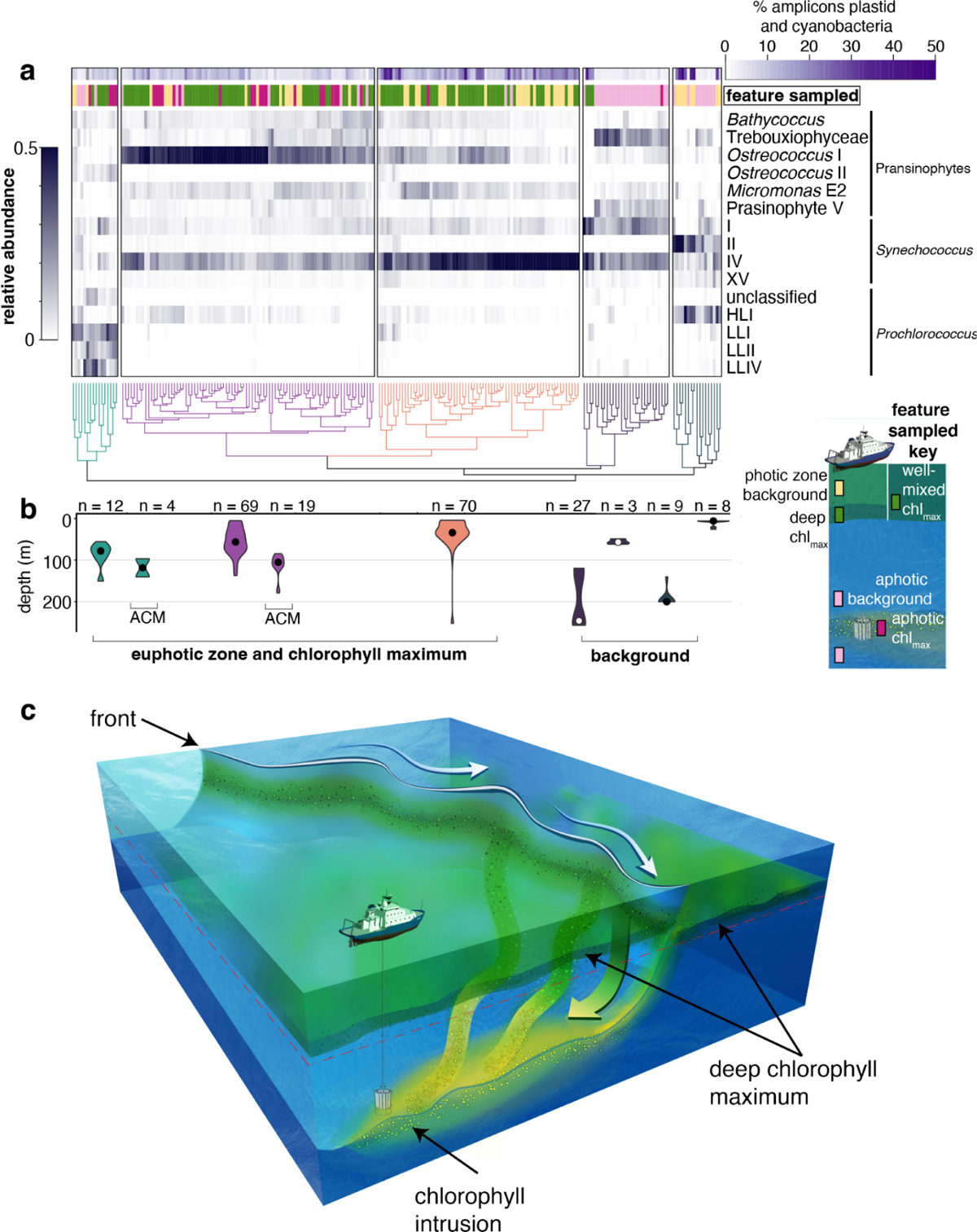
Surface ocean picophytoplankton communities are exported to the aphotic zone via intrusions. **a,** Hierarchical clustering of picophytoplankton taxa identified using phylogenetic placement of V1-V2 16S rRNA gene ASVs (see Extended Data Fig. 8 for phytoplankton taxa of lesser relative abundance). Each colored cluster in dendrogram is significant (*p<*0.05). First, we tested the extent to which diversity had been saturated in our deep sequencing efforts, analysis of rare taxa was not undertaken due to lack of saturation in some samples (Extended Data Fig. 9). The top horizontal bar (greyscale) indicates the percentage of ASVs from cyanobacteria and photosynthetic eukaryotes out of the total number of amplicons, which includes sequences from heterotrophic bacteria; the two left hand most clusters show extremely low percentages of sequences from phytoplankton, as expected due to being classical dark ocean samples. The bar above the heatmap indicates the sampled feature type. **b,** respective depth distribution of samples in selected clusters (vertical boxes, and above clustering) represented as the mean and distribution of samples (n indicates number of samples). **c,** Schematic of key features in a frontal system with a deep chlorophyll maximum and an aphotic chlorophyll-containing intrusion. Intrusions are coherent, 3-dimensional features with multiple origin locations along the front. Sampling occurred in transects capturing a cross section of a 3-dimensional system.

The depth distributions of samples in each cluster demonstrate that ACM communities are significantly deeper in the water column than communities within the same cluster (30–50 m deeper than the depth of origin, *p<*10^−5^, Fig. 2b). The limited change in community composition, as well as physical parameters, between photic zone and intrusion samples suggests that advective transport to 100-200 m occurs over a few days (*<*10days). Our high-resolution comparative analyses of picophytoplankton community composition served as a sensitive tracer of along-isopycnal vertical transport and connectivity between the ACM and source water communities. The results further establish that intrusions drive export of picophytoplankton from photic zone chlorophyll maxima operating on a year-round basis, rather than just during physically energetic periods of the year, like winter.

### POC transported within intrusions into the upper mesopelagic includes a dominant signal from non-photosynthetic bacteria

The intrusions transported picophytoplankton to the dark ocean, raising questions regarding the total flux of carbon to the upper mesopelagic from intrusions. Using beam transmission data as a proxy for POC, we found that all instances of high POC in the aphotic zone at the observed fronts were associated with ACMs that result from intrusions, rather than sinking (Fig. 3a,b). The intrusions generated substantial patchiness in POC in the upper mesopelagic. This was most evident in the probability density distribution of POC differentiated into ‘intrusion’ and ‘background’ mesopelagic samples with the same depth distribution. The total POC within intrusions averages twice that of background mesopelagic waters (Fig. 3a,b).

**Fig. 3.**
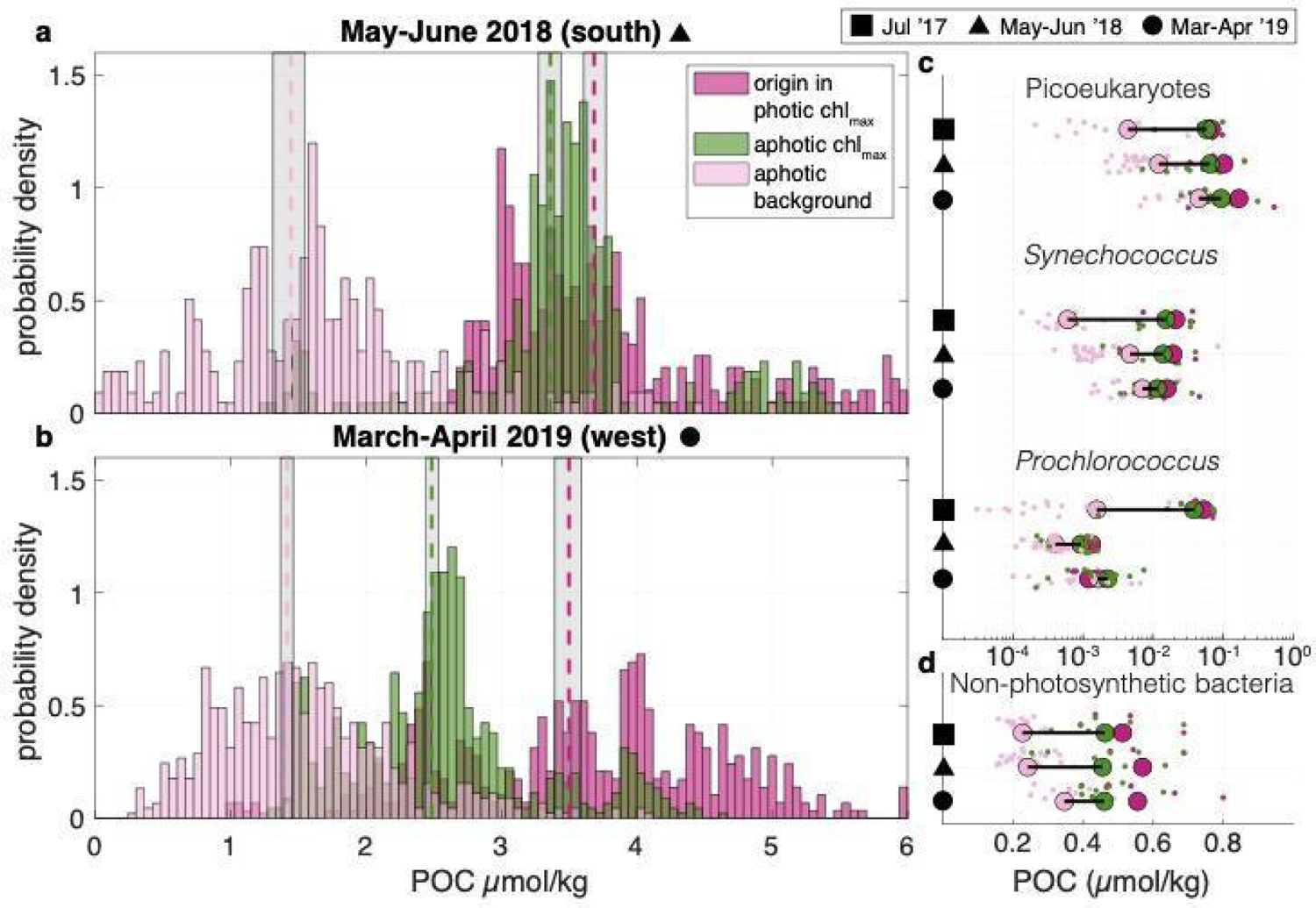
Intrusions are enhanced in particulate organic carbon at depth with unexpectedly large contributions from non-photosynthetic bacteria. **a, b,** Probability density functions of POC concentration estimated for summer 2018 and spring 2019, respectively, from transmissometer profiles within the ACM (dark pink), background (light pink, randomly sampled from outside the ACM to have the same depth distribution), and origin waters (green, high chlorophyll regions in the photic zone with density ranges of ACM). The geometric mean of POC distributions in the ACM, origin, and background are indicated (dashed lines, color coding on panel) with gray shading for the 95% confidence interval (note, beam transmission and optical backscatter are lacking for 2017). **c,** POC amount contributed by each major phytoplankton group over the three expeditions (y-axis black symbols). Note log scale (X-axis). **d,** POC contributed by non-photosynthetic bacteria is greater than phytoplankton. For **c, d** POC estimates are based on flow cytometry analysis of intact cells and established conversion factors; colors are as in panel A. Note that improved data on taxon-specific cellular carbon content will be an important future step for improving quantitative estimates of taxon-specific carbon export.

The adaptive sampling of intrusions enabled us to quantify contributions of picophytoplankton groups to physically-driven export for the first time. The contribution of *Prochlorococcus*, *Synechococcus* and picoeukaryotic phytoplankton to carbon export is assessed using cell abundances and established carbon conversion factors. This reveals that across all seasons studied, there is a significant enhancement of carbon from picophytoplanktonic cells in aphotic regions of the intrusions compared with background mesopelagic waters at the same depth, where very few picophytoplankton are observed (Fig. 3c). Picophytoplankton biomass in the aphotic intrusions is of comparable magnitude and composition to the photic zone communities.

Because the intrusion export mechanism also carries heterotrophic microbes that co-occur with the picophytoplankton from the photic zone, we enumerated them by flow cytometry. The process of subduction might be important for heterotrophic bacteria, since they are even smaller than picophytoplankton and sink more slowly. Non-photosynthetic microbes averaged 870,000 ± 320,000 cells ml^−1^ in the photic zone and, due to their high abundance, comprised about half of the total microbial POC in sunlit waters (Fig. 3c,d), akin to data from prior subtropical studies^31^. Subduction of these abundant bacteria resulted in abundance peaks in intrusions (560,000 ± 150,000 cells ml^−1^) that were nearly twice the number seen in background upper mesopelagic samples (310,000 ± 78,000 cells ml^−1^) at the same depth in the aphotic zone (Fig. 3d). Moreover, their biomass contributed as much, or more, POC to advective export as all the picophytoplankton groups combined (Fig. 3c,d).

The substantial POC contribution by heterotrophic bacteria was initially surprising, since the intrusions herein were identified via their elevated chlorophyll. Indeed, the presence of anomalously high chlorophyll and backscattering below the photic zone^12,13^ has been considered almost exclusively through the lens of phytoplankton and the importance of phytoplanktonic advective export in the biological pump. However, our findings are consistent with the fact that in the nutrient-limited subtropics non-photosynthetic bacteria have much higher abundances in the photic zone – where we demonstrate intrusions originate – than at depth^32^. Thus, the intrusion export mechanism represents an ‘optimal’ export for heterotrophic bacteria, matching their peak biomass. Moreover, reevaluation of the relationship between total POC (beam transmissometer-derived) and microbial POC (cell counts-derived) showed that when biomass of picophytoplankton is summed with that of heterotrophic bacteria, there is a significant correlation with total POC (R^2^ = 0.42 (spring), 0.46 (summer), *p*<10^-14^), stronger than with just phytoplankton (R^2^ = 0.39 (spring), 0.42 (summer), *p*<10^-11^), with seasonally-dependent offsets connected to amounts of POC from intact cells versus detrital particles (Extended Data Fig. 7). The discovery of a marked enrichment of carbon in the upper mesopelagic due to heterotrophic bacterial communities that are transported within 3D-intrusions modifies current concepts of microbial carbon export.

### Enigmatic non-photosynthetic bacterial groups are stimulated in intrusions

Differences in the physiological and biogeochemical impact of transport into the dark ocean along intrusions are notable for heterotrophic bacteria in comparison with picophytoplankton. The latter are at life’s end, providing important labile food resources for predators and bacteria, and potentially for export to greater depths. In contrast, some non-photosynthetic bacteria have the possibility to continue growing at depth, in addition to having differentiated roles in remineralization of POC and attenuation of organic matter^24,32^. Thus, along with the large export contributions in terms of carbon, the impacts on biogeochemical transformations and ecological trajectories of transport along intrusions are perhaps greatest for heterotrophic microbes.

Subtropical non-photosynthetic bacterial communities are dominated by SAR11 in the photic zone^24^. Here, we observed dominance by SAR11 clade Ia in the photic zone while SAR11 clades Ib and II and SAR324 had the highest relative abundances in aphotic communities (Fig. 4a). Heterotrophic microbial communities reflected the surface origin communities in a manner that differed from the phytoplankton communities. Specifically, the intrusions increased the aphotic zone heterotrophic community complexity and multiple non-photosynthetic bacterial taxa appeared to systematically respond to subduction in differing ways (Fig. 4a). Differential relative abundance analysis on the ASV data, revealed 16 non-photosynthetic bacterioplankton ASVs with significant increases in the ACMs over both aphotic background waters and intrusion origin waters. Among these were SAR406 (8 ASVs), the *Verrucomicrobia* lineages Arctic 97B-4 (4 ASVs) and MB11C04 (marine groups) (1 ASV), and OCS116 clade members (3 ASVs). The response of *Verrucomicrobia* and OCS116 to labile material from phytoplankton blooms has been noted previously^19,33,34^, supporting the idea that they preferentially respond to the decay of the relatively high phytoplankton biomass in intrusions. SAR406 have also been observed to respond to environmental variability at mid-depth in other regions, but with less clear triggers^21^. This broad uncultivated alphaproteobacterial lineage exhibits diverse roles in sulfur and nitrogen metabolism, based on metagenomic analyses, with some clades even encoding nitrous oxide reductase. Thus, the biogeochemical transformations they perform within the intrusions appear complex and likely shift the repertoire of the collective metabolisms present in the aphotic zone.

**Fig. 4.**
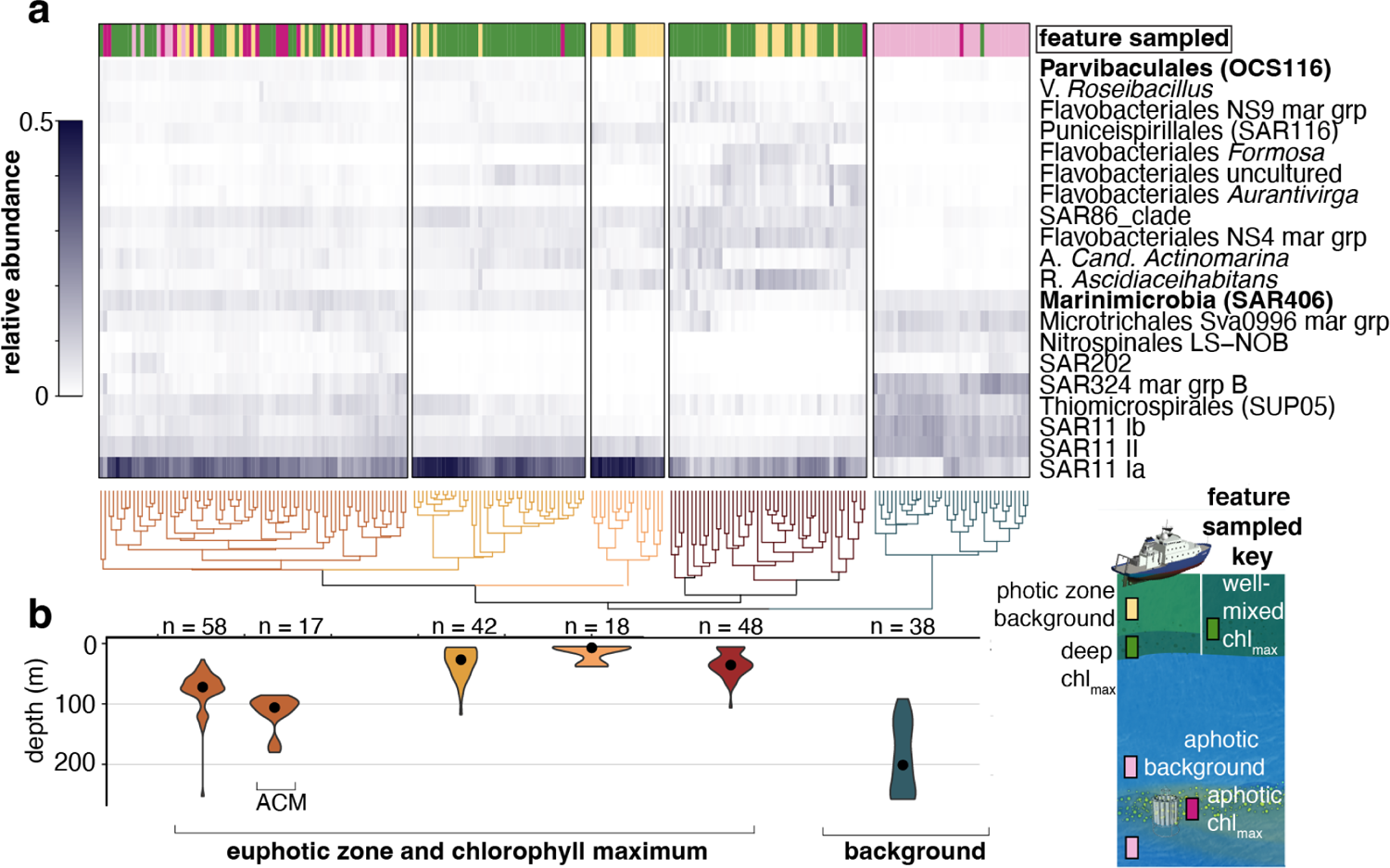
Specific non-photosynthetic bacterial lineages appear to respond to intrusion conditions, reshaping community structure. **a,** Hierarchical clustering of non-photosynthetic bacterioplankton distributions with taxonomy determined using V1-V2 16S rRNA gene ASVs and established databases. The percentage contribution to total amplicons of the most relatively abundant taxa across all samples is shown in the heat map (grey scale). Bacterial taxa that wee differentially more abundant in intrusions based on statistical analysis are denoted (bold lettering) and indicate enrichment in the ACM relative to the upper mesopelagic background and source photic-zone waters. The sample feature is shown in the bar above the heatmap and is subtly different than that of picophytoplankton due to differences in hierarchical cluster influenced by enrichment of heterotrophs that appear to respond to the intrusion conditions. Each colored cluster is significant (*p<*0.05). **b,** respective depth distribution of samples in selected clusters (vertical boxes, and above clustering) represented as the mean and distribution of samples (n indicates number of samples). Nearly all ACM samples are in the same cluster and ACM samples are found deeper than other communities in this cluster.

Subduction through intrusions, as described herein, impacts the functional and metabolic diversity of the aphotic zone differently than processes that are conventionally invoked for explaining vertical connectivity (e.g. convection, active transport by zooplankton, and sinking detrital particles^35–37^). The more rapid timescales of intrusions, and coherence of the water mass and environmental conditions, contribute to the living nature of exported cells and growth of activated community members as they experience transitions in light and organic matter lability during transport.

### Vertical flux of POC due to intrusions can rival sinking flux

We next aimed to estimate POC contributions to the dark ocean via intrusions by combining multiple datasets and process study modeling for the frontal setting in a global context. Using the observed POC anomalies in intrusions and a conservative estimate of the vertical velocity as 10 m day^−1^, we estimate the local flux in Western Mediterranean Sea intrusions is on the order of 100 mg C m^−2^ day^−1^.

Subduction promotes two-way connectivity between photic and aphotic communities. Therefore, to obtain the net carbon flux, it is necessary to integrate in space and time over regions and episodes of positive and negative flux (i.e. upwelling and downwelling). We used a submesoscale resolving process study ocean model initialized with the observed hydrographic conditions and POC distribution (Extended Data Text 9). Certain aspects of the large-scale Mediterranean circulation are particular, resulting in quasi-stationary fronts that have previously been thought to be critical for generating intrusions^16,18,22^. The process study model we use here generalizes the frontal export process beyond the quasi-steady topographically-influenced conditions in the Alborán Sea and instead quantifies the influence of globally-relevant submesoscale and mesoscale baroclinic instability. Due to the submesoscale nature of the observed dynamics, downwelling (subduction) occurs more rapidly and in a smaller spatial area than upwelling, and water remains subducted for many days. As as result, there is net subduction of carbon through an episodic flux on the order 100 mg C m^−2^ day^−1^, as observed, and an integrated downward advective flux of 10 mg C m^−2^ day^−1^ at a depth of 150 m (Extended Data Fig. 11). Sinking POC flux estimates from time series observations in the North Atlantic and Western Mediterranean range from 0.3–60 mg C m^−2^ day^−1^.^38,39^ The achieved advective export flux is of similar magnitude to the average sinking flux and, notably, its inclusion in coarse resolution carbon cycle models or budgets doubles the rate of ocean carbon export.

Intrusions occur along density surfaces and the mechanism hinges on mesoscale processes, which set up deep-reaching fronts and increase the potential depth range of export, and episodic submesoscale processes, which accelerate the rate of export and disrupt periodic flows. Together, these increase the likelihood that water parcels are not upwelled after subduction. Furthermore, rather than persisting at the ACM density surface, modeling efforts indicate that POC can be transported below the depth of downward vertical motion, when sinking and advective export occur together^40^. Future work should quantify the ways that the depth structure and timescale of vertical transport interacts with ecological dynamics and sinking fluxes.

We next assessed the global extent of POC export by intrusions using physical understanding of their dynamics and a number of data sources, including the Bermuda Atlantic Time-series, which characterizes the subtropical ocean water column from the surface to mesopelagic on a monthly basis^41^, satellite observations, and the global array of ARGO floats^42^. Specifically, along-isopycnal transport was estimated using a well-established parameterization of eddy tracer flux^43^ and a diffusivity estimate based on recent work that accounts for submesoscale dynamics^44^, alongside the global observations (Extended Data Text 8). We quantify flux as

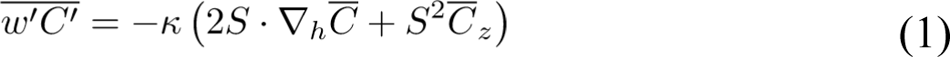

where *w*^′^*C*^′^ is the spatial and temporal average vertical flux of POC (*C*) and *S* is the isopycnal slope. The eddy transfer coefficient is given by 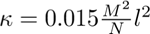 where *M*^2^ and *N*^2^ are the lateral and vertical buoyancy gradients, respectively, and *l* is a mixing length (Extended Data Fig. 10). This is a conservative estimate of carbon flux because it underestimates the influence of submesoscale processes. Several patterns emerged, including a large flux of POC by intrusions in subpolar gyres, which act in concert with the seasonal mixed layer eddy-driven flux^12^. The advective flux of POC in the seasonally stratified subtropical gyres, at ∼75 mg C m^−2^ d^−1^ and 50 mg C m^−2^ d^−1^ in northern and southern subtropical gyres, respectively, is larger than previous estimates. Moreover, in 26 years of BATS data, 1.3% of profiles have an ACM that is likely due to intrusions from the photic zone (Extended Data Fig. 13), lending support to our process model-based estimates. These results establish the global magnitude of the intrusion export process, however total contributions to POC export will be better constrained with the future acquisition of subsurface POC data alongside high-resolution modeling studies.

Our analyses indicate that the net intrusion-driven POC flux at 100 m is downwards and is of similar magnitude as the sinking flux in subtropical regions^45^ (Fig. 5). Current estimates of POC export in subtropical oceans are known to be insufficient when it comes to accounting for heterotrophic production levels in the mesopelagic, or for balancing observed rates of net community production in the photic zone on annual timescales^46,47^. Other physically-driven processes including mixed layer eddy subduction^12^ and the mixed layer pump^48,49^ do not resolve this imbalance on annual timescales. It should be noted that some global and regional modeling studies also conclude that additional processes beyond sinking flux are needed to close the carbon budget in subtropical systems^50,51^. In addition to the enhanced POC flux, intrusions carry dissolved organic matter thereby contributing to a larger and chemically-complex carbon pool at depth^52^. Organic carbon export by intrusions is likely critical to closing carbon budgets in subtropical regions.

**Fig. 5.**
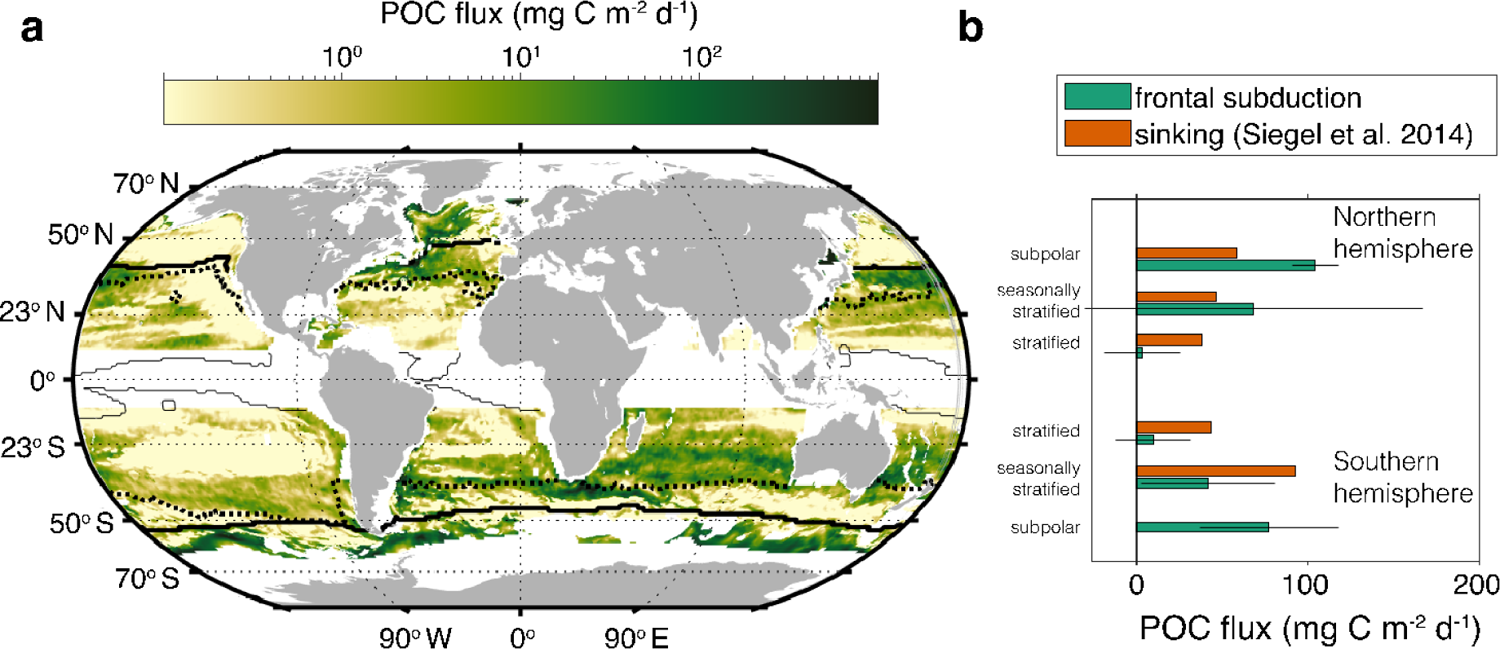
Global importance of intrusion driven particulate organic carbon export. **a,** Global estimate of the magnitude of POC flux due to physically-driven intrusions from the base of the photic zone in subtropical and subpolar gyres. The thick line delineates the equatorward limit of the subpolar gyre, the thick dotted line divides the seasonally stratified subtropical gyre from the permanently stratified subtropical gyre, and the thin solid line delineates the equatorward limit of the subtropical gyres. **b,** Average POC flux from physically-driven intrusions grouped into three broad bioregions in each hemisphere (subpolar, seasonally stratified subtropical [seasonally stratified], and permanently stratified subtropical [stratified]). Error bars on the integrated POC flux represent errors in estimated POC gradients, which can be large due to relatively sparse observations. The subduction flux is compared to the sinking flux at the base of the photic zone as estimated by Siegel et al.^45^. There is no error estimate provided for the sinking flux.

## Conclusions

The coupled ocean biophysical dynamics revealed here transport picophytoplankton and heterotrophic bacteria communities in three-dimensional intrusions from photosynthetically productive sunlit layers to the mesopelagic in subtropical oceans. This constitutes an important contribution to POC export and microbial diversity in the upper mesopelagic of the Mediterranean, and is relevant globally. The export mechanism operates throughout the year and recruits contributions from the biomass maximum of the photic zone, including the DCM, which was not thought to contribute substantially to export. Many of the small cell size populations that are exported by intrusion dynamics are unlikely to have other routes into the mesopelagic as intact cells. By revealing this process, we highlight how the ecology and biogeochemistry of the ocean’s interior are influenced by surface-intensified dynamics that are variable on timescales of days to weeks rather than only seasonal timescales.

Transport of photic zone communities to depth in intrusions alters the composition of food supplies and contributes greater functional complexity to mesopelagic ecosystems. Heterotrophic bacteria constitute a large fraction of the microbial POC in intrusions and injection of these communities is ecologically significant because some taxa appear to thrive in the aphotic zone thereby modifying ‘standard’ mesopelagic microbial metabolisms and interactions. Moreover, the oxygen transported within intrusions alongside the dissolved and particulate organic carbon will impact the biogeochemistry of the dark ocean, including remineralization and attenuation of organic matter. There is much to be learned about the uncultivated bacteria that are enhanced in the intrusions. As the oceans warm and density stratification intensifies, picophytoplankton are expected to constitute a larger fraction of primary production^6,7^. The physically-driven export mechanism described here is predicted to be robust to climate change impacts on increasing photic zone stratification and related selection of smaller sized taxa. Overall, we show that intrusions are ubiquitous in subtropical oceans, and are responsible for the coupling of photic-zone microbes and production with mesopelagic productivity, biodiversity, and biogeochemical processes.

## Materials and methods

### Sampling strategy

Sampling occurred during three research cruises that took place from July 17-24, 2017 on the *R/V SOCIB* (IRENE), May 27-June 2, 2018 on the *NRV Alliance* (Calypso 2018; CLP18) and March 21-April 12, 2019 on the *NO Pourquoi Pas?* (Calypso 2019; CLP19)^53^. Ship transects were oriented perpendicular to the largest buoyancy and velocity gradients to the extent possible. This sampling design aims to capture the greatest degree of variability in water masses and isopycnal depth range. Each transect was completed as rapidly as possible. On transects where CTD casts were performed to sample water (Fig. 1a) there were 30 minutes to 1 hour between casts on a given transect. 3-8 CTD casts were conducted in each cross-frontal transect reported here, at a spacing of 1–10 km. These transects were completed in 3-6 hours. CTD cast locations were chosen based on cross-front underway CTD (UCTD) or EcoCTD^54^ measurements performed in the same cross-front section immediately prior to the CTD transect. Samples were collected at 6 depths using Niskin bottles mounted to a CTD rosette. On every cast, samples were collected (i) near the surface at a fixed depth of 5m; (ii) within the mixed layer (∼25 meters); (iii) the chlorophyll maximum, i.e. either the DCM or the base of the mixed layer; (iv) just below the chlorophyll maximum; (v) the secondary, aphotic chlorophyll maximum (ACM) if present, or a fixed depth of 120 meters if not present; (vi) the maximum cast depth (200 m in 2017 and 2019, 250 m in 2018). On 3 of the transects, community composition data is not available but we use nutrient, CTD, and bio-optical observations.

### Hydrographic observations

Temperature and salinity were measured with a dual Seabird SBE9 on the CTD rosette. Near continuous profiling was carried out with an *Oceansciences* underway-CTD or the EcoCTD. The UCTD, a custom SBE sensor that samples at 16 Hz. The EcoCTD measures conductivity, temperature, and pressure with an RBR*concerto*^3^ inductive CTD^54^. The data processing from the underway instruments is described in Dever et al. ^55^. Discrete samples were taken to calibrate the SBE9. The UCTD was calibrated immediately prior to the cruise by Seabird. The EcoCTD was calibrated by RBR immediately prior to the cruise and the optical instruments were cross-calibrated by mounting it to the CTD rosette frame for one cast during each cruise. Velocity was observed using an acoustic Doppler current profiler (ADCP) mounted to the vessel on each cruise.

### Bio-optical observations

Chlorophyll *in vivo* fluorescence was measured by a Chelsea Aqua 3 fluorometer (CLP18) and WETLabs ECO-AFL/FL (CLP19) mounted on the CTD and a WETLabs ECOPuck on the EcoCTD. *In vivo* fluorescence was calibrated with *in situ* samples collected from the fluorescence maximum and at the maximum profile depth (200 or 250 m). For each sample, 0.5 L of seawater was filtered onboard through a 45 mm GF/F Whatman filter. Concentrations of chl *a* were determined using the fluorometric method described in^56^, total pigment extracted with 90% acetone for 24 hours at 4^◦^C in the dark and determined with a fluorometer (Turner Designs).

Oxygen was measured by a Seabird SBE43 oxygen sensor mounted to the CTD and a RINKO III oxygen sensor on the EcoCTD. The output of the oxygen sensors was calibrated against the Winkler titrations^57^ performed at sea within 48 hours of sampling (Metrohm 888 Titrator).

### Biological community composition

#### DNA Extraction, PCR, and Sequencing

DNA samples for characterizing microbial communities were obtained by filtering biomass from 500 ml of natural seawater onto 47 mm 0.2 *µ*m pore size polyethersulfone membrane filters (Supor 200, Pall Gelman). Filters were placed into sterile cryovials, flash-frozen in liquid nitrogen and transferred to −80^◦^C until extraction using the DNeasy Plant Kit (Qiagen), with a modification including a bead beating step^58^. During the 2017 and 2018 cruises, samples were stored in liquid nitrogen for the duration of the cruise after which they were stored at −80^◦^C until extraction. DNA was amplified using the primers 27FB (5^′^-AGRGTTYGATYMTGGCTCAG3^′^) and 338RPL (5^′^-GCWGCCWCCCGTAGGWGT-3^′^)^59,60^ targeting the V1-V2 hypervariable region of the 16S rRNA gene with Illumina adapters, allowing us to explore the diversity of eukaryotic phytoplankton (via plastid 16S rRNA gene), cyanobacteria, and heterotrophic bacterioplankton^28,59–61^. PCR reactions contained 25 ng of template, 5 *µ*l of 10× buffer, 1 U of HiFi-Taq, 1.6 mM MgSO4 (Thermo Fisher) and 0.2 *µ*M of each primer. The PCR cycling parameters were 94^◦^C for 2 min; 30×94^◦^C for 15 s, 55^◦^C for 30 s, 68^◦^C for 1 min, and a final elongation at 68^◦^C for 7 min. Paired-end library sequencing (2 × 300bp) was performed using the Illumina MiSeq platform.

#### 16S rRNA gene amplicon analyses

Sequences were demultiplexed and assigned to corresponding samples using CASAVA (Illumina). A 10 bp running window was utilized to trim low-quality sequence ends at a Phred quality (Q) of 25 using Sickle 1.33^62^. Paired-end reads were merged using USEARCH v10.0.240^63^ when reads had a ≥ 50 bp overlap with maximum 5% mismatch. The merged reads were then filtered to remove reads with maximum error rate *>* 0.001 or shorter than 200 bp. Only sequences with exact match to both primers were kept and primer sequences were trimmed using Cutadapt v.1.13^64^. After removal of single sequence reads, the average number of amplicon reads from the 317 samples was 259,104±133,685. Cyanobacterial and plastid amplicons were initially parsed using the phylogenetic pipeline in PhyloAssigner v.6.166^59^. Amplicons from plastids and cyanobacteria were further classified using a global plastid and cyanobacterial reference alignment and tree according to protocols outlined in Choi et al.^61^. Cyanobacterial amplicons were analyzed using fine-scale cyanobacterial reference alignment and tree^60^, and plastid amplicons were further analyzed using multiple high-resolution alignments and trees focusing on stramenopiles and viridiplantae^28,61^.

For the assessment of heterotrophic bacterioplankton communities, sequences (82,135,874 reads) were resolved into 49,150 amplicon sequence variants (ASVs) of 100% pairwise identity with USEARCH v10.0.240^63^. Taxonomies were assigned to each ASV using classify-sklearn by QIIME2^65^ searching against the SILVA database release 138^66^.

#### Cell quantification

We preserved samples for quantification by flow cytometry from the same Niskin bottles from which we took the samples for DNA extraction. Samples were preserved using 10 *µ*l of EM grade 25% Glutaraldehyde per 1 ml seawater and then flash frozen in liquid nitrogen (or at −80^◦^ for casts 1-19 in 2019). Samples were then stored at −80^◦^C until analysis. Cells were quantified using a BD Influx flow cytometer equipped with a 488 nm laser. Forward angle light scatter, side scatter, and autofluorescence at 692/40 nm, 572/27 nm, and 520/35 nm were recorded. Photosynthetic cells were counted by triggering on forward angle light scatter. Calibration beads were added to each sample immediately before analysis (0.75 *µ*m yellow-green, Polysciences, Inc and 1.0-1.4 *µ*m ultrarainbow, Spherotech). Each sample was run for 8 minutes at 25 *µl* min^−1^ after a pre-run of 2 minutes. Heterotrophic bacteria cells were counted by staining samples with SYBR Green and triggering on autofluorescence at 520 nm. Samples were run for 4 minutes after a 2-minute pre-run. Analysis was performed in WinList (Verity Software House).

Integrating the constrained gene copy numbers in cyanobacteria to examine specific clade abundances^60^ allowed us to leverage flow cytometric data to examine contributions of specific *Prochlorococcus* and *Synechococcus* clades (this analysis was not performed for picoeukaryotes due to having unknown rRNA gene copy numbers).

### Statistical analysis of community composition

The hierarchical clustering presented in the main text was performed using the “ggdendro” package and average clustering and Manhattan distance. In all cases, intrusion samples refer to the aphotic portion of intrusions, unless otherwise specified. This is defined as deeper than 90 meters, using the maximum 1% light level across all observations.

We test which ASVs have statistically significant different abundances (differentially abundant) between sample groups using ANCOM implemented in Qiime2. We define three groups for this analysis: the chlorophyll maximum on the dense side of the front and intrusion samples shallower than 90 m (origin waters), the samples in the aphotic zone in intrusions (deeper than 90 meters), and samples in the background deeper than 90 m. ANCOM assumes that fewer than 25% of the ASVs change between groups^67^.

### Estimating particulate organic carbon (POC)

We estimate the POC concentration using multiple complementary methods. Each captures a different contribution to the POC.

#### Flow cytometry

We use literature values^68,69^ to estimate the carbon content of the cells enumerated by flow cytometry. The values we use are 39, 82, and 530 20 fg C cell^−1^ for *Prochlorococcus, Synechococcus,* and picoeukaryotes, respectively. The carbon per cell is most uncertain for the picoeukaryotes, which have the largest biomass per cell and are a large contributor to the small photosynthetic community during the 2019 cruise. Following (*22*) we use a value of 20 fg C cell^−1^ for heterotrophic bacteria^70^.

#### Optical instruments

The particle beam attenuation is a proxy for particles in the size range 5-20 *µ*m (*24*). The beam transmission (Tr) (Seatech/Chelsea in 2018, WETlabs in 2019) was measured and used to derive *c_p_* as

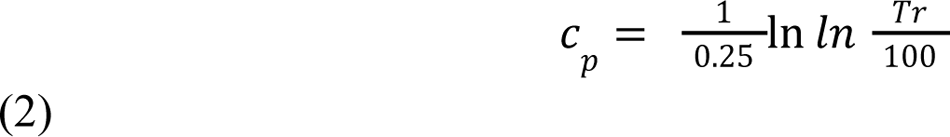

The community composition, in addition to the total POC, may affect *c_p_*. The beam attenuation is affected by phytoplankton and also by bacteria, zooplankton, and detritus. It is not currently possible to measure beam transmission on the EcoCTD. Instead, the EcoCTD measures backscatter at 700 nm. Both the beam attenuation and backscatter can be converted to carbon using region-specific empirical calibrations. We use a correlation between *c_p_*and POC of 1.78 m^2^(g C)^−1^ ^71^. The range of possible values used for uncertainty estimates includes the values reported in^72^. We use a relationship between backscatter and POC (POC = 54463 × *b_bp_* − 1.19 mg C m−3)^73^.

### Definition of intrusion ACMs

Intrusions are identified as an aphotic chlorophyll maximum (ACM).

1. The ACM must extend at least 5 m in the vertical. The bounds of the intrusion are set by the bounds of the elevated chlorophyll concentration. *There are thermohaline intrusions that are not associated with elevated chlorophyll concentration. These are intrusions in a more general sense but are not included in the analysis here because they do not necessarily contribute more to carbon export than the background waters*.
2. Intrusions are present continuously in more than one CTD cast within a transect. The intrusion must be below 100 m in at least one cast along the transect. Being present continuously means that the intrusion is present within the same density range across the front. *This criterion allows us to confidently identify thin intrusions (small vertical extent) over large lateral scale. In addition, we are able to approximate the origins of the intrusions within the euphotic zone by identifying the intrusions when they are above 100 m. By contrast, when identifying intrusions from isolated profiles*^74^ *investigators were only able to identify intrusions that were already below 100 m and that were at least 150 m in vertical extent*.
3. The ACM is associated with an inversion in the vertical in at least one other property (temperature, salinity, oxygen, POC). *This criterion helps to ensure that the ACM is due to an intrusion rather than local growth or sinking of phytoplankton. Within an intrusion, the gradients in chlorophyll and the other properties may not be aligned due to patchiness in the origin location*.

### Process study model

The POC flux is computed using process study models initialized with a cross section across the Almería-Oran front. Two configurations of the Process Study Ocean Model (PSOM)^75,76^ are run with one representing a stratified summer period and the other the springtime period when the thermocline density surface outcrop at the sea surface^26^. The initial condition representing the springtime period is obtained by cooling the sea surface and using convective adjustment until the mixed layer has a maximum depth of 70 m. The models are re-entrant channels with walls in the north-south direction and periodic boundaries in the east-west direction. The model domain is centered at 36.9°N and extends 128 km in the east-west direction and 216 km in the north-south direction and is 1 km deep. The models have grid spacing of 500 meters with a stretched grid in the north-south direction that attains a spacing of 2 km within 40 km of the southern and northern solid boundaries. There are 64 vertical levels on a stretched grid with grid spacing ranging from 0.5 m at the surface to 54 m at depth. The model timestep is 108 s. The horizontal diffusion is 1 m^2^/s. The vertical diffusion has a constant value of 10^−5^ m^2^/s. The model has a flat bottom and a linear bottom drag of 10^−4^ m/s. The POC flux is computed by evolving the mean POC profile from each season for a month with no reactions or restoring.

### Eddy flux parameterization

To approximate the global magnitude of POC export due to the intrusion process described in this paper, we parameterize the vertical flux using a skew flux^43,77^

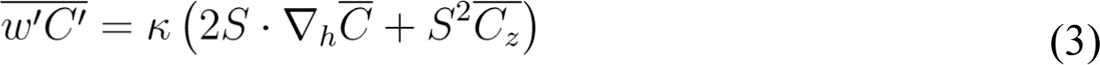

where *w* is the vertical velocity, *C* is the POC concentration, and *S* is the isopycnal slope. The isopycnal slope is a 2D vector and is computed from a monthly Argo climatology^78^. This climatology has a resolution of 56 km near the equator and increases to 22.5 km at 66.5° and poleward. The isopycnal slope term in this expression accounts for the effect of a limited depth range of sloping isopycnal surfaces. The flux will go to zero if an isopycnal is flat. We use an eddy transfer coefficient given by 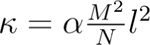. This eddy transfer coefficient was derived by Visbeck et al.^77^ based on linear stability analysis of baroclinic eddies. The transfer coefficient, which has units of m^2^s^−^1, is given by an eddy velocity multiplied by a mixing length (*l*). The eddy velocity is determined to be *αRi*^−1*/*2^*fl*. It is notable that the coefficient *αRi*^−1*/*2^ is the scaling derived by Freilich and Mahadevan^44^ for the proportion of the vertical motion that is along sloping isopycnal surfaces. This observation further supports the use of this scaling for the vertical eddy flux. We use the constant *α* = 0.015^77^. The coefficient 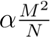 is computed from a monthly climatology of Argo profiles^78,79^. We quantify the mixing length from the variance of salinity on an isopycnal surface as 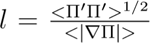 where Π = Π – <u>Π</u> after Cole et al.^80^. All gradients and the mixing length are computed on a 1-degree grid using Argo data. A single annual value is used for the mixing length and monthly values are used for 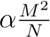. The average values are shown in Extended Data Fig. 10.

The POC gradients are computed from a climatology of POC based on Biogeochemical Argo data and satellite observations combined with a neural network to create 3D fields and 0.25° resolution with 36 vertical levels between the surface and 1000 meters^81^. The POC flux is averaged over all data points where the density difference between the surface and the euphotic depth is at least 0.3 kg m^−3^, indicating that the base of the euphotic zone is stratified.

## Acknowledgements

This research is a part of the CALYPSO Departmental Research Initiative (DRI) funded by the U.S. Office of Naval Research (ONR). We are grateful to the captains and crews of the *B/O SOCIB*, *NRV Alliance*, and *N/O Pourquoi Pas?*. Funding was also provided by a NDSEG Fellowship, Martin Fellowship, Grassle Fellowship and the Montrym fund to MAF, the Gordon and Betty Moore Foundation (GBMF 3788) and NSF Dimensions 2230811 to AZW. We are grateful to Pierre Chabert, Kausalya Mahadevan, Alex Beyer, Sebastian Essink, Margaret Conley, HM Aravind, and Salvador Veira for assistance with sample collection and David Needham, Charmaine Yung, Eugenio Cutolo, Craig Carlson, and Stephen Giovannoni for thoughtful discussions.

## Data availability

Hydrographic data displayed in the figures will be available at Zenodo DOI (XX) and sequences with all metadata will be available upon publication at BioSample accession number XX.

## Author contributions

MAF, AM, and AZW conceptualized the study; MAF, AM, MD, EA, JA, AC, SR, AP, JTF, TMSJ, and ED executed the field sampling; MAF, CP, LS, and CJC performed analysis of biological samples; MAF and MD calibrated hydrographic observations; JTF and TMSJ collected underway hydrographic observations; EA, JA, and AC calibrated biogeochemical observations; AM and MAF conceptualized the modeling; MAF performed the formal data analysis, performed the modeling studies, and wrote the original draft; AM supervised; all authors reviewed and edited the manuscript.

## Notes

### Competing Interest Statement

The authors have declared no competing interest.

