## Supplementary Information for "3D-intrusions transport active surface microbial assemblages to the dark ocean"

### Extended Data
### 3D intrusions actively transport surface microbial assemblages to the dark ocean

#### 1 Water mass structure

There are three typical water masses in the Alborán Sea, which are characterized by their temperature and salinity properties on a given density surface. The Atlantic water is relatively fresh and cool, the Mediterranean water is relatively warm and salty and the Modified Atlantic Water lies between those two water masses in temperature-salinity space [1]. All three water masses were sampled during the field campaigns presented in this manuscript. The sampling during IRENE (July 2017) was in the Mediterranean water and Atlantic water. During the other two field campaigns, sampling was in the Modified Atlantic Water and the Atlantic water (Extended Data Fig. 1). During the stratified sampling periods (July 2017 - IRENE and May-June 2018 - CLP18), the surface water was much warmer than the interior, inhibiting water mass exchange.

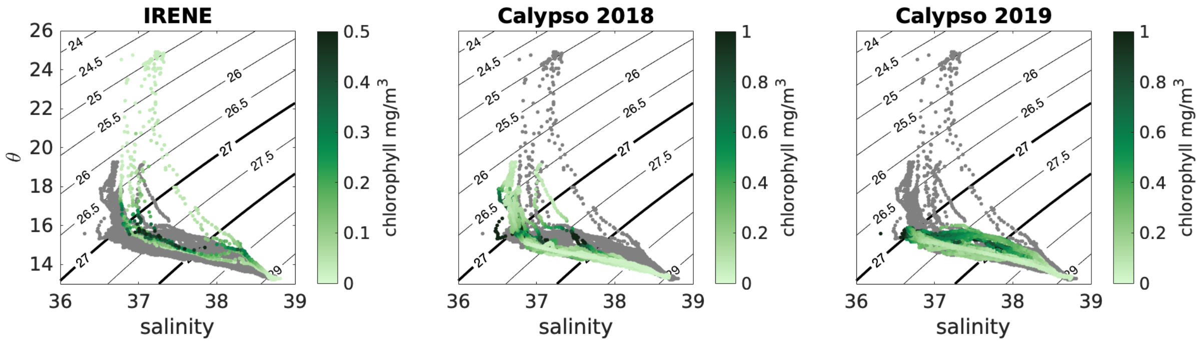


**Extended Data Figure 1**: Temperature-salinity diagrams showing the water mass structure in the Alborán Sea. Contours are density. The gray points are repeated on each panel and show the water mass structure sampled by CTD casts over the three field campaigns. Chlorophyll concentration is plotted in temperature-salinity space for each field campaign. The branches in temperature salinity space show the water masses. The chlorophyll maximum is on a different density surface in each water mass, reflecting the variable nutrient-density relationship.

#### 2 Depth distribution of biomass

Advective export occurs due to the co-occurrence of biomass and downward vertical water mass transport. Therefore, the depth distribution of both the community and the vertical velocity play a key role in

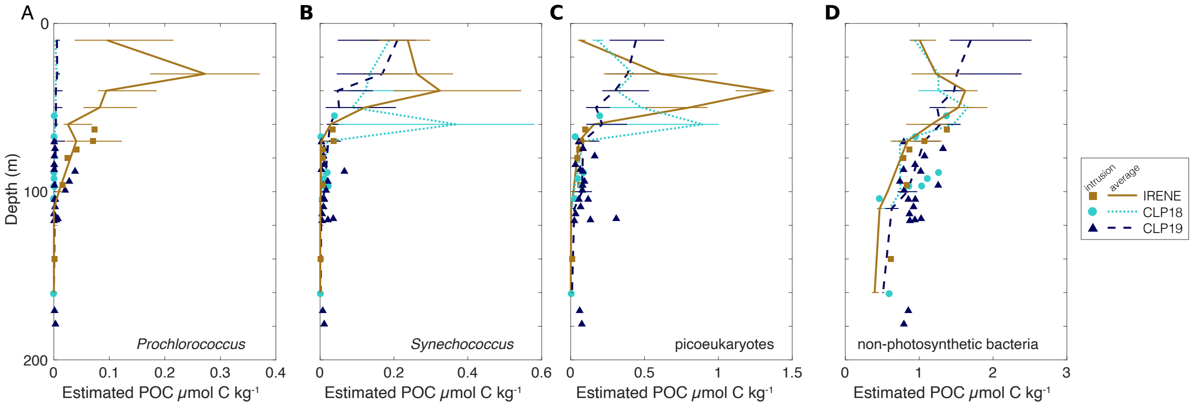


**Extended Data Figure 2**: Average profiles of plankton estimated by enumerating cells with flow cytometry and using taxa-specific cell to biomass conversions. The solid line shows the median in depth bins, the error bars show the interquartile range. The points indicate the concentration of each group in the intrusions. The samples are grouped by research cruise.

determining the total flux and the community composition of the flux. The average profiles of cyanobacteria, picoeukaryotes, and non-photosynthetic bacteria differ between the research cruises, which represent regional and seasonal variability (Extended Data Figure 2). During the July 2017 research cruise in the oligotrophic Mediterranean water, *Prochlorococcus* was abundant and had peak biomass in the Atlantic water mass DCM near 40 m depth. Low-light *Prochlorococcus* also peaked in the Mediterranean water DCM near 70 m depth. By contrast, *Prochlorococcus* was in low abundance during the spring and early summer. Instead, *Synechococcus* and picoeukaryotes had high abundance in the chlorophyll maximum layer, which was in the surface in March-April 2019 and deeper during May-June 2018 and July 2017. Note that the maximum photosynthetic biomass was on average deeper during May-June 2018 at around 60 m than in July 2017. However, during the research cruise in July 2017, the DCM on the Atlantic water mass side of the front was much shallower and had higher POC, which dominates the average profile, than on the Mediterranean side of the front, which is the source of the intrusion communities.

The biomass and chlorophyll maxima were coincident in the observed profiles (Extended Data Fig. 3), as has been reported previously in this region [2].

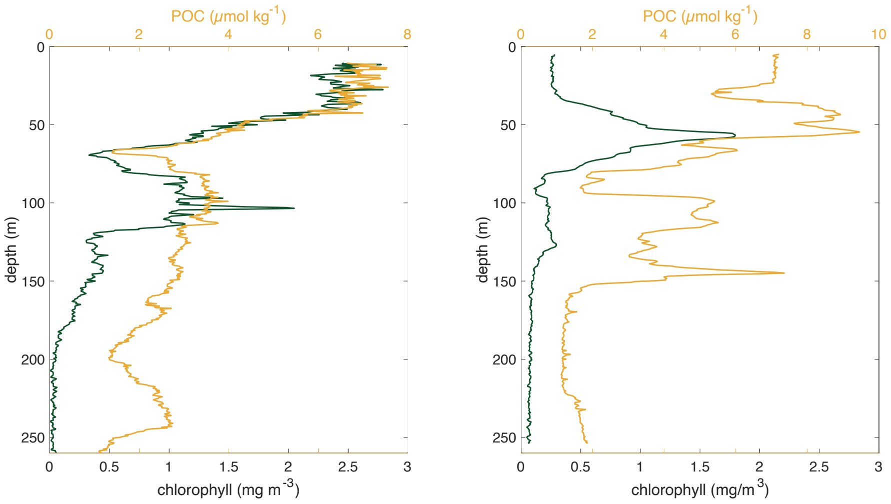


**Extended Data Figure 3**: Chlorophyll and POC profiles for the same profiles shown in main text Figure 1b,c demonstrate that chlorophyll and POC have coincident maxima. In this figure, POC is using observations from a transmissometer.

### 3 Cell fluorescence

We find limited deviations in average chlorophyll *a* fluorescence per cell measured from flow cytometry in ACMs compared with similar water masses from the photic zone, suggesting that there is limited photoacclimation and that the enhanced chlorophyll is not explained by photoacclimation (Fig. S7). We measure the average red (692 nm) and orange (572 nm) fluorescence of each taxonomic group using the flow cytometer. While a few of the intrusion samples that are observed, particularly *Synechococcus* cells display an increase in the average cell fluorescence, beyond that of any shallower or deeper communities, most of the intrusion samples that are observed have cell fluorescence that is consistent with the surrounding communities, or the communities that are found shallower than the intrusion samples (Extended Data Fig. 4).

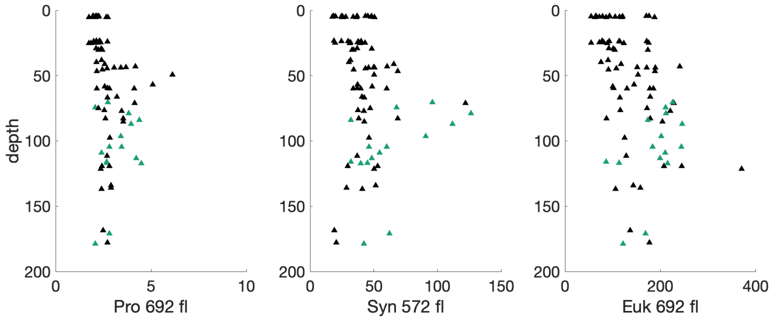


**Extended Data Figure 4**: Average fluorescence of the cells observed during CLP19 normalized by yellow-green and rainbow beads in each taxonomic group, *Prochlorococcus*, *Synechococcus*, and picoeukaryotes as a function of depth. The *Prochlorococcus* and picoeukaryotes fluorescence is chlorophyll-a fluorescence while the *Synechococcus* fluorescence is phycoerythrin fluorescence. The green symbols show samples from intrusions.

### 4 Case study of early spring physical export

The transects observed during March–April 2019 demonstrate that physically-driven export into the pycnocline takes place throughout the year, even when most of the photosynthetic biomass is in the surface mixed layer. This not only reveals that frontal dynamics can export carbon throughout the year, but also that water mass transport can occur from the base of the mixed layer to below the depth of the winter mixed layer [3]. While we observed that the community composition was not as strongly differentiated between the Atlantic Water and Modified Atlantic Water as it was between the Atlantic water and Mediterranean water, we were still able to observe anomalous biological communities at depth. These communities were anomalous in both the total abundance of plankton and the composition of the biological community relative to the surrounding water at the same depth. What is particularly notable is that the inversions in AOU and temperature are associated with an inversion in biomass. For example, Extended Data Fig. 5 at 53 km shows that the sample within the intrusion had higher biomass, but was deeper than a sample outside the intrusion. In addition, the intrusion has a high abundance of high light *Prochlorococcus*, which is not expected to grow at the depths below 100 m where it was observed. By contrast, in the denser water outside of the intrusion a low light *Prochlorococcus* ecotype is observed.

On the transect plotted in Extended Data Fig. 5 we measured the photosystem II efficiency of the cells (measured as Fv/Fm) and found that it decreases from 0.42 at 50 m to 0.29 at 75 m to 0.19 at 100 m even as the phytoplankton community composition changes little, suggesting that the phytoplankton cells are stressed [4]. The photosystem II efficiency (Fv/Fm) of algae was assessed by examining the changes in chlorophyll fluorescence with the electron transport inhibitor 3.(3,4-dichlorophenyl)-1,1-dimethylurea (DCMU) that blocks electron transport at the electron acceptor Q in PSII [5]. Minimum and maximum fluorescence were measured after 30 min of dark adaptation using a Turner Designs 10-AU fluorometer on board. Studying the changes in the photosynthetic community composition along intrusions may reveal mortality processes.

### 5 Biogeochemical contrasts in intrusions

The intrusions have significantly higher POC and lower AOU than the background. The magnitude of this difference varies between the research cruises likely due to differences in the ecology and biogeochemistry regionally and over time (Table 1).

| location | POC | | | AOU | | |
|---|---|---|---|---|---|---|
| | intrusions | background | difference | intrusions | background | difference |
| CLP19 | [2.35, 2.44] | [1.39, 1.48] | 0.9 | [41.9, 43.4] | [46.1, 48.9] | 4.3 |
| CLP18 | [3.27, 3.45] | [1.30, 1.52] | 2 | [56.9, 59.2] | [64.1, 67.8] | 8.4 |

**Table 1**: POC concentration ($\mu$mol/kg) and AOU ($\mu$mol/kg) in intrusions below 100 m and outside the intrusions ("sampled background"). The intervals show the bootstrapped 95% confidence interval of the geometric mean of each category (1000 iterations). The concentration outside the intrusions is calculated by averaging the POC and AOU concentration from a random sample of points with the sample depth distribution as the intrusion samples. The variation in the geometric mean and the confidence intervals among random samples from the background is less than 0.001 $\mu$mol/kg. The location is the research cruise and region. CLP19 WAG is transects C1–5. CLP18 is transects B1 and B2. The t-test p value that the geometric mean in the intrusions is significantly different from the sampled background is less than 0.001 in all cases.

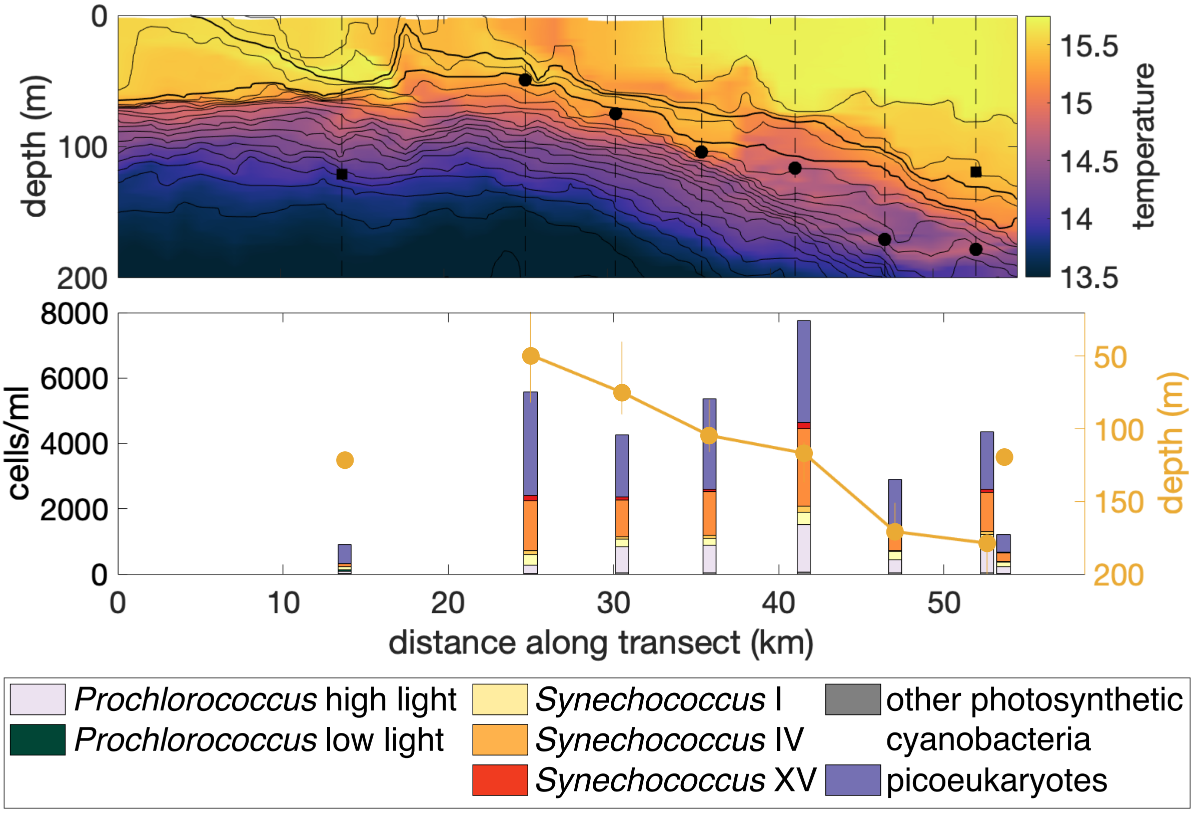


**Extended Data Figure 5**: (upper panel) Example 2D section across the western Alborán Gyre (transect C2). Temperature measured by the UCTD is shown. The locations of CTD casts are shown in dashed lines. Selected water samples are shown to compare and contrast the community composition within and outside the intrusion on this transect. The cast at 36 km is shown in main text Fig. 1B. (lower panel) Community composition at selected sampling locations, combining cell counts from flow cytometry and 16S rRNA gene amplicon sequencing. The depth of the samples is shown by the orange points. The intrusion samples are circles and are connected by a solid line.

### 6 Comparison between methods for estimating POC

#### 6.1 Backscatter and beam transmission

In the northwestern Mediterranean, the diel cycles of $c_p$ and $b_{bp}(700)$ have been observed to show only weak correlation indicating that there are differential contributions of phytoplankton to these proxies [6]. $c_p$ shows larger diel variations [7] suggesting that small detritus and bacteria make larger contributions to $b_{bp}$ than to $c_p$. At calibration casts where the EcoCTD is attached to the ship-board CTD frame, we find a linear correlation between $c_p$ and $b_{bp}(700)$ (Extended Data Fig. 6).

#### 6.2 Flow cytometry and beam transmission

A comparison of the quantification of POC from flow cytometry and the transmissometer reveals good agreement and useful differences (Extended Data Fig. 7). There is a strong correlation between POC estimated from flow cytometry and POC estimated using the transmissometer. This indicates that the

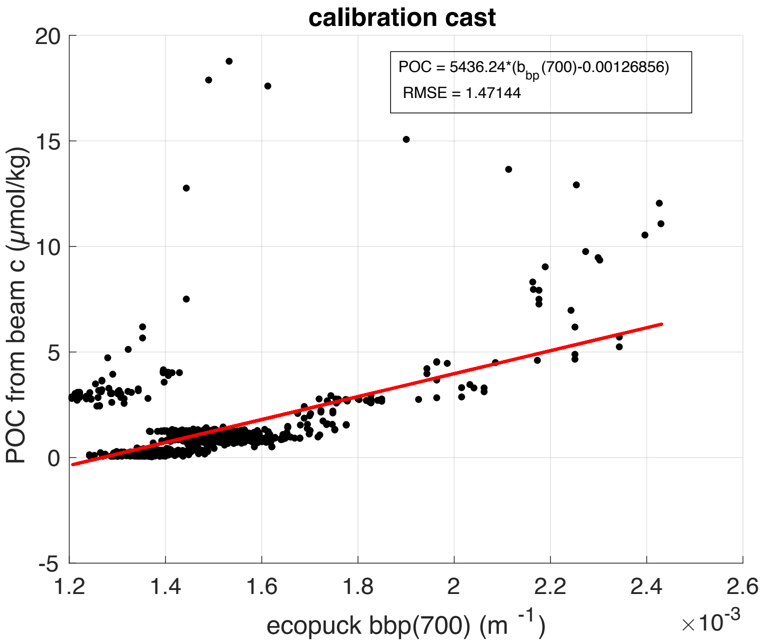


**Extended Data Figure 6**: Correlation between POC estimated from the transmissometer mounted to the CTD rosette and from bbp(700) measured by the EcoCTD during a calibration cast where the EcoCTD was attached to the CTD rosette frame.

microbial communities play a key role in the variability of POC. However, the transmissometer estimates are approximately twice as large as the estimates from flow cytometry during CLP19 with a larger discrepancy during CLP18. The slope in log-log space is 1:1 for the CLP19 samples, steeper than 1:1 for CLP18 transect 1 samples, and shallower than 1:1 for CLP18 transect 2 samples. Where the slope is 1:1, we can infer that the additional biomass captured by the transmissometer is likely detritus and heterotrophic eukaryotes. Where the slope is not 1:1, we may infer that the biomass conversions have systematic errors. In the case of the estimates from flow cytometry, this may indicate that the eukaryotes are smaller (shallower slope than 1:1) or larger (steeper slope than 1:1) than estimated for the conversion from cell counts to biomass (see section Eukaryotic community composition). Alternatively, the transmissometer may systematically over (steeper) or under (shallower) estimate the POC concentration at high concentrations due to patchiness in the community composition.

Interpreting results from flow cytometry alone allows us to focus on the impacts of physical processes on the export of small intact cells while use of the transmissometer reveals the net impact on POC export, including detritus and large organisms that may settle more quickly or perform diel vertical migration.

#### 6.3 Eukaryotic community composition

There is more variability in the cell size and cellular carbon content of eukaryotes than bacteria. This makes estimation of the POC from cell enumeration more uncertain when the communities are dominated by eukaryotes. Furthermore, we focus on the role of the intrusion process in the export of small cells. Therefore, it is important to verify that the intrusion communities are composed of small cells, even when eukaroytes are numerically dominant.

In agreement with the general trend of increasing cell abundance with decreasing cell size, we find that samples with a large number of eukaryotes are dominated by relatively small cells, as estimated

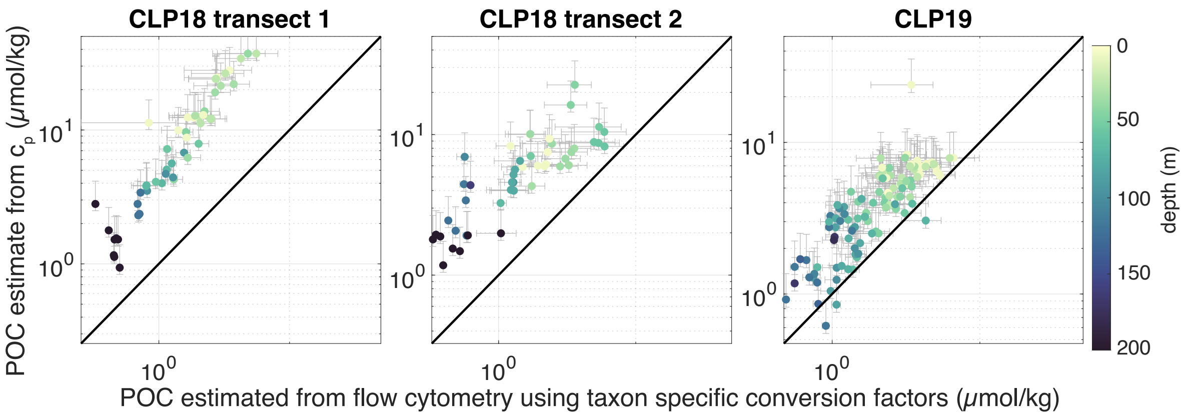


**Extended Data Figure 7**: Comparison of estimates of POC concentration from flow cytometry (FCM) and transmissometer ($c_p$). Transects 1 and 2 from CLP18 (transects B1 and B2, respectively) are plotted separately because they display different slopes. All samples from CLP19 are plotted together. IRENE is not shown because there is no estimate of POC from a transmissometer during IRENE. The error bars show uncertainty due to conversion factors from beam transmission and cell counts to POC.

by FALS (forward angle light scattering), and Viridiplantae amplicons, which tend to be smaller than Stramenopiles, the other dominant group identified from plastids (Extended Data Fig. 8). The variation in average FALS between 200 and 400 results in a factor of 2.5 difference in estimates of carbon per cell assuming a cell size conversion of $\frac{\text{size cell}}{\text{size beads}} = \frac{\text{FALS cells}}{\text{FALS beads}}^{x}$ where $x$ is between 4 and 6 and log C (pg/cell) = 0.94× log Vol (μm$^{-3}$)-0.6 [8].

### 7 Microbial diversity

Rarefaction curves are used to examine the influence of sequencing depth on the estimation of the microbial community composition. The lack of saturation of the curves indicates the sequencing depth precludes analysis of rare taxa (Extended Data Fig. 9).

### 8 Eddy flux parameterization

To approximate the global magnitude of POC export due to the intrusion process described in this paper, we parameterize the vertical flux using a skew flux [9, 10]

$$\overline{w'C'} = \kappa \left(2S \cdot \nabla_h \overline{C} + S^2 \overline{C_z}\right) \tag{1}$$

where $w$ is the vertical velocity, $C$ is the POC concentration, and $S$ is the isopycnal slope. The isopycnal slope is a 2D vector and is computed from a monthly Argo climatology [11]. The isopycnal slope term in this expression accounts for the effect of a limited depth range of sloping isopycnal surfaces. The flux will go to zero if an isopycnal is flat. We use an eddy transfer coefficient given by $\kappa = \alpha \frac{M^2}{N} l^2$. This eddy transfer coefficient was derived by Visbeck et al. [10] based on linear stability analysis of baroclinic eddies. The transfer coefficient, which has units of m$^2$s$^-$1, is given by an eddy velocity multiplied by a mixing length ($l$). The eddy velocity is determined to be $\alpha Ri^{-1/2} fl$. It is notable that the coefficient $\alpha Ri^{-1/2}$ is the scaling derived by Freilich and Mahadevan [12] for the proportion of the vertical motion that is along sloping isopycnal surfaces. This observation further supports the use of this scaling for the vertical eddy flux. We use the constant $\alpha = 0.015$ as in [10]. The coefficient $\alpha \frac{M^2}{N}$ is computed from a monthly

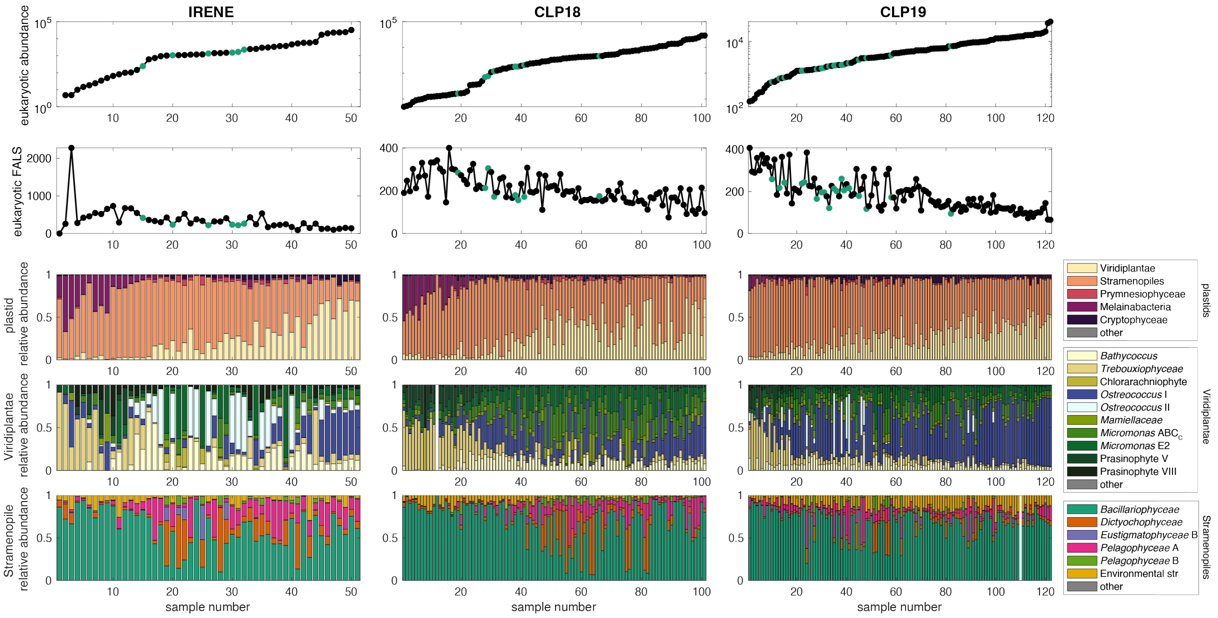


**Extended Data Figure 8**: The photosynthetic eukaryote abundance and cell size show systematic variation with the community composition. Samples are arranged in order of ascending abundance of picoeukaryotes, as enumerated by flow cytometry. Each column is a different field experiment. The first row shows eukaryote abundance. The second row shows geometric mean forward angle light scatter (FALS), a proxy for cell size. The green dots are intrusion samples. The third to fifth rows show the relative abundance of amplicons in three taxonomic groups: third row, plastids without cyanobacteria; fourth row, Viridiplantae; fifth row, Stramenopiles.

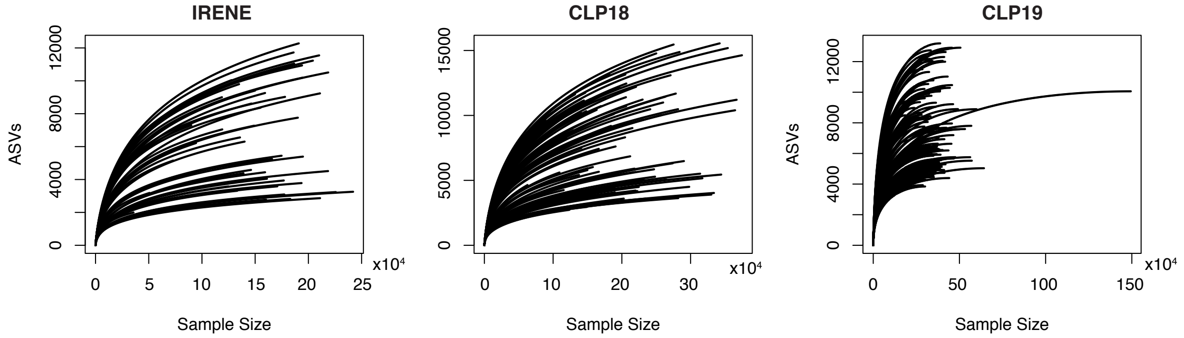


**Extended Data Figure 9**: Rarefaction curves for the three sampling campaigns

climatology of Argo profiles [11, 13]. We quantify the mixing length from the variance of salinity on an isopycnal surface as $l = \frac{<\Pi'\Pi'>^{1/2}}{<|\nabla\Pi|>}$ where $\Pi' = \Pi - \overline{\Pi}$ after Cole et al. 2015 [14]. All gradients and the mixing length are computed on a 1-degree grid using Argo data. A single annual value is used for the mixing length and monthly values are used for $\alpha\frac{M^2}{N}$. The average values are shown in Extended Data Fig. 10.

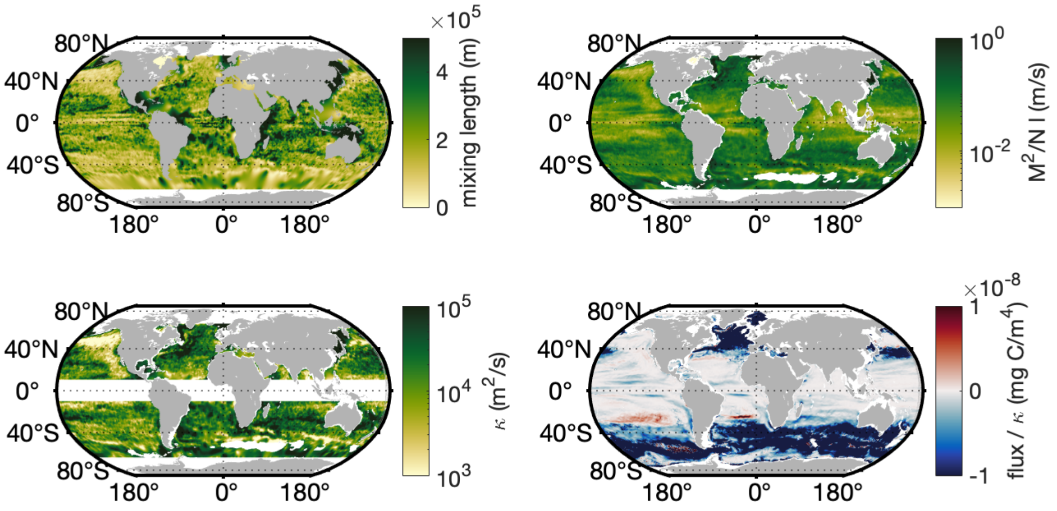


**Extended Data Figure 10**: Components of POC flux calculation

Coverage of observations of direct measurements of POC and optical proxies of POC are limited globally, resulting in a relatively high error in the POC magnitude and POC gradients compared to the other quantities. While the results presented here suggest that POC flux from the base of the euphotic zone could be important, the global generalization is limited by the data availability. The global distribution of optical backscatter observations used in training the algorithm is shown in figure 4 of Sauzede et al. 2021 and displays a bias in observation density in the Southern Ocean, Mediterranean Sea, and North Atlantic and very few observations in the low latitude subtropics [15]. It is imperative to expand observations of subsurface biological communities.

This parameterization for the vertical flux is based on mid-latitude baroclinic instability and is limited in its application to equatorial regions, where parameterization based on the density gradients does not capture the full scaling relationship in Freilich and Mahadevan 2019 [12] and where the eddy velocity inferred by $\alpha Ri^{-1/2}$ is small while the eddy kinetic energy in models is large. Therefore, these regions are excluded from the analysis. This parameterization can only be applied below the mixed layer and parameterizes mesoscale and large submesoscale processes. The parameterization differs from the one used by Omand et al. [16], which only parameterizes the impact of mixed layer eddies.

### 9 Modeling frontal dynamics

#### 9.1 POC flux

The POC flux averaged over a frontal region is computed using process study models initialized with hydrographic structure from observations in the Western Mediterranean Sea. Two models are run with one representing the stratified period (when the initial condition was sampled) and the other the springtime period when the thermocline density surface outcrop at the sea surface [17]. The initial condition representing the springtime period is obtained by cooling the sea surface and using convective adjustment until the mixed layer has a maximum depth of 70 m. The has an idealized configuration with no topographic features. It is a re-entrant channel configuration with closed boundaries in the north and south and periodic boundaries in the east-west direction. The model horizontal resolution is 500 m, except near the closed north and south walls where the cell length increases linearly to 2 km. The model is evolved with a horizontal diffusivity of 1 $m^2s^{-1}$ and a vertical diffusivity of $10^{-5}$ $m^2s^{-1}$. The POC flux is computed by evolving the mean POC profile from each season for a month with no reactions or restoring. The total POC flux is estimated as the average carbon concentration integrated between the model base and 90 m (Extended Data Fig. 11).

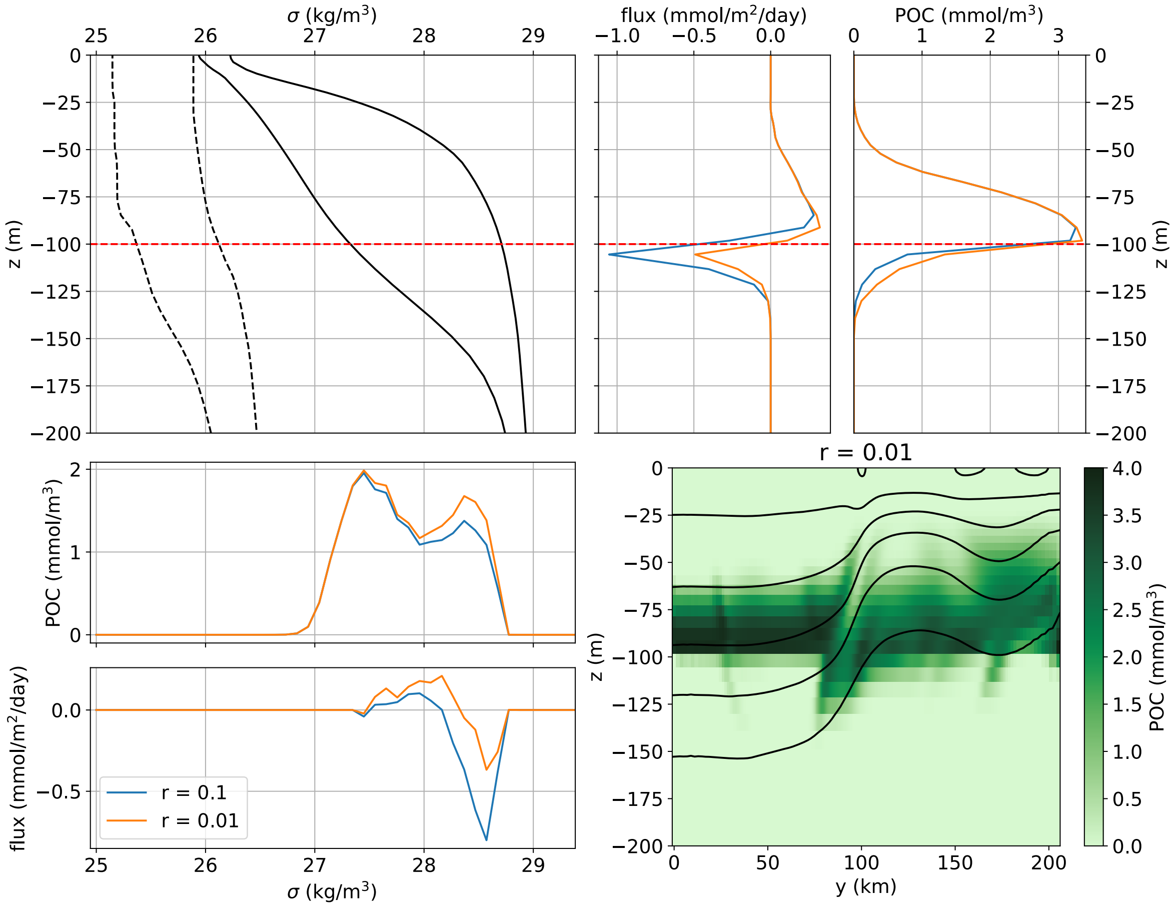


**Extended Data Figure 11**: Model POC flux. The flux, computed as $wP$ where $P$ is the POC concentration and $w$ is the vertical velocity, is shown as a function to depth (upper right) and density (lower left). An example model cross section is shown in the lower right for comparison to observations. The upper left panel shows the density range at each depth that are used for model initial conditions. The results in this figure and Extended Data Fig. 12A-C are from the model with solid lines (stratified period) while the results in Extended Data Fig. 12D are from the model with dashed lines.

#### 9.2 3D structure of POC intrusions

While intrusions are identified in cross-frontal transects in the present work, they are three-dimensional features that are advected along the front. The shape of the intrusions evolve as they are advected. In a two-dimensional slice an intrusion may appear across a wide density range [18]. In the case of the features identified here, because the intrusions originate from a frontal region where, by definition, there is a

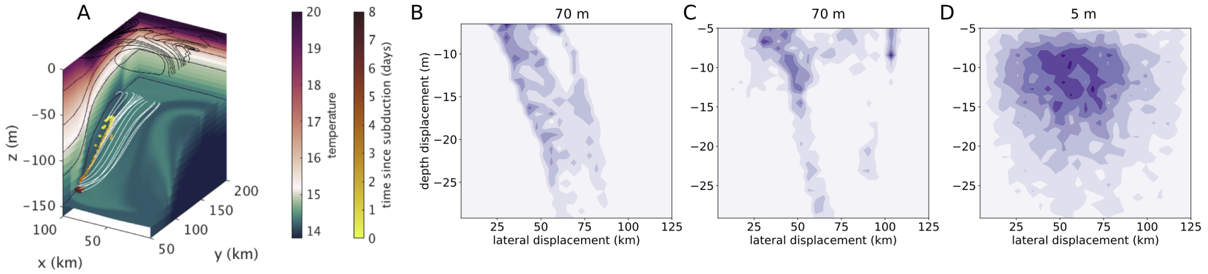


**Extended Data Figure 12**: Model analysis of subduction. (A) Lagrangian particles (yellow) originated at 70 m and are subducted along an isopycnal surface. The white trajectories show the x-y-z location history of the particles. The isopycnal surface is shown with temperature shading. Temperature is shown on the boundaries of the frame as well. (B-D) Histograms of the relationship between vertical and lateral displacement of water parcels. (B) Particles initially at 70 m in a model with a stratified surface layer representing the June and July research cruises. The particles shown in panel (A) are a subset of the particles shown in panel B. (C) Particles initially at 70 m in a model with a 50 m deep mixed layer representing the March-April research cruise. (D) Particles initially at 5 m in the model with a 50 m deep mixed layer. Statistics on this plot are from water parcels that leave the mixed layer.

large range of density over the relatively small origin location, we expect intrusions occur across a wide range density. Furthermore, we find that there is not an exact correspondence between biogeochemical and thermohaline intrusions. This is because biogeochemical and thermohaline gradients differ in both the vertical and horizontal.

We examine the shape of subducted features in the model, which allows us to propose 3D structures of POC intrusions and examine their temporal evolution. Due to the along-front current, water parcels are moving more quickly along-front than either downwards or across the front. Model analysis shows that water parcels come from 25–100 km upstream of the location where they are observed subsurface (Extended Data Fig. 12). In two process study models with the same thermocline density structure but distinct near surface stratification (one stratified as in July 2017 and one with 50-70 m deep mixed layers as in March-April 2019), there is a positive relationship between lateral and vertical distance on water parcels that subduct within the thermocline that corresponds approximately to the isopycnal slope in the thermocline (Extended Data Fig. 12B,C). Water parcels that are deeper have traveled farther from their origin location both in the horizontal and vertical with the ratio of those two motions set by the isopycnal slope. In these models, the water parcels that originate within the thermocline experience subduction along a well-defined isopycnal surface and their motions show the signature of the mesoscale meander [19]. The relationship between vertical and lateral motions is less coherent for water parcels that subduct from the mixed layer where the isopycnal slope is less well-defined and there is a greater influence of small-scale processes (Extended Data Fig. 12D; [3]).

### 10 Observations at the Bermuda Atlantic Time Series

Given the limitations in estimating the POC flux globally from observations, we demonstrate the relevance of this export process in other regions by observing that ACMs are ubiquitous in the subtropical gyres. For example, at the Bermuda Atlantic Time Series ACMs are present below 200 meters in over 5% of the monthly profiles in the late spring (Extended Data Fig. 13).

While the density of the mixed layer varies throughout the year, the density of the chlorophyll maximum layer is approximately constant at BATS from the onset of stratification in March to April until the mixed layer deepens (and becomes denser) in the fall. During the stratified period, ACMs are generated from the base of the DCM, as indicated by the ACMs lying within the DCM density range, but with a slightly higher average seawater density. There is a distinct seasonality to ACM occurrence, with ACMs occurring more frequently in the late spring and early summer, before the mixed layer begins to deepen.

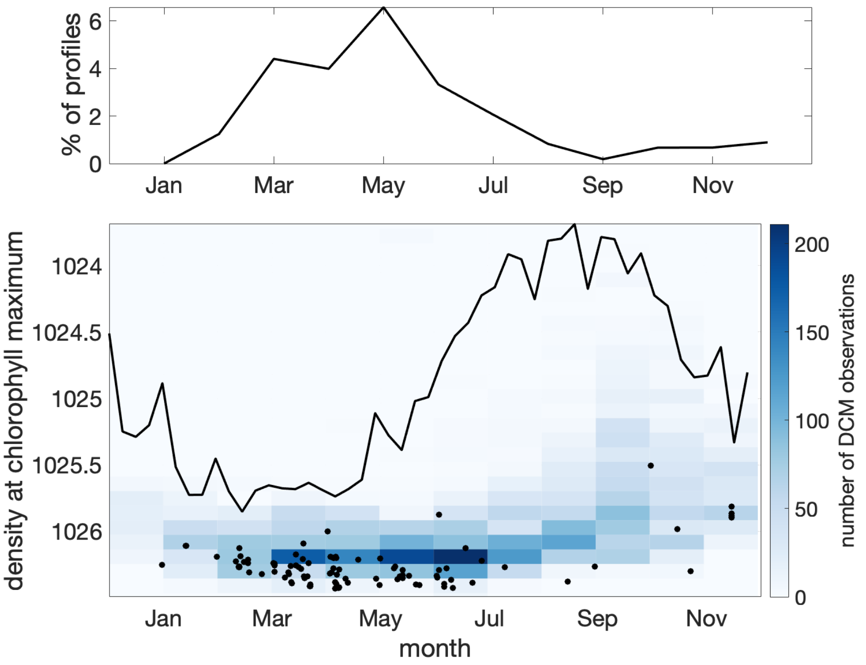


**Extended Data Figure 13**: Time series of observations of aphotic chlorophyll maxima at the Bermuda Atlantic Time Series (BATS). The top panel shows the percentage of monthly profiles in which a chlorophyll maximum is observed below 200 meters. The lower panel shows the observations in density space. The background color shows the frequency with which the DCM occurs on a given density surface in a given month. The black line shows the mixed layer density. The black dots show the time and seawater density of aphotic chlorophyll maximum observations.

### 11 Depth range of subduction

Export by intrusions has a distinct vertical profile from sinking flux (Extended Data Fig. 10). Advective subduction is constrained to occur along sloping density surfaces. This means that the maximum depth to which subduction occurs is determined by the density structure of the region where subduction occurs. While our observations are from fronts concentrated in the upper 200 m, in the open ocean, eddy effects can extend below 1000 m [20] and density surfaces that outcrop in the photic zone can extend downwards hundreds of meters (Extended Data Fig. 13). We introduce the concept of "potential displacement" to quantify the depth influence of subduction. Subduction can occur along a given isopycnal surface only within the depth range of that isopycnal surface. In depth-density plots, this depth range can be diagnosed. For example, near BATS water parcels on the $\sigma$ =26.25 surface can subduct 300 m from 100 m to 400 m and water parcels on the $\sigma$ =26.35 surface can subduct 450 m from 100 m to 550 m (Extended Data Fig. 14). The density variation at a given depth, particularly in the upper ocean, reflects both large scale gradients and eddy dynamics. In this region near BATS, the density variations are due to eddy dynamics and do not display large scale gradients. While potential displacement can be quantified from hydrography alone, the actual displacement depends on the ageostrophic velocities. The depth over which

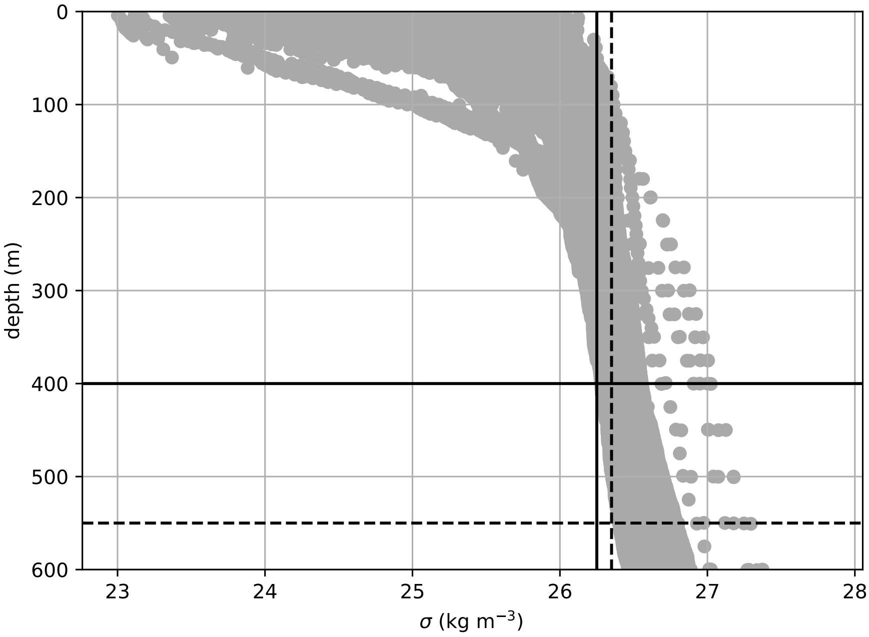


**Extended Data Figure 14**: Density as a function of depth from Argo profiles from May to August in a 3-degree box surrounding the Bermuda Atlantic Time Series station. The black lines show the depth range of the 26.25 (solid) and 26.35 (dashed) density surfaces.

advective export fluxes attenuate depends not just on the remineralization rate, as it does for sinking flux, but also on the vertical structure of the vertical motion.

Carbon may be exported deeper than would be possible by subduction along through a combination of subduction and sinking or mixing [21].

#### 12 Flow cytometry gating strategy

Figures exemplifying the gating strategy for photosynthetic and heterotrophic microbes are shown in Extended Data Figures 15 and 16.

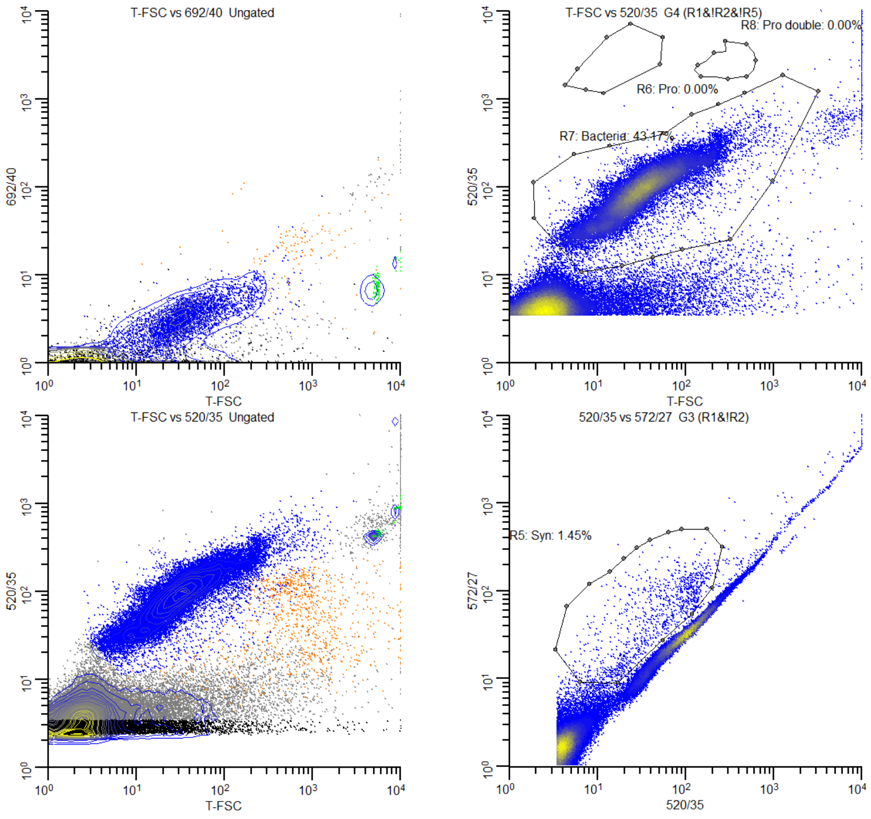


Id1: MAF_4Sep18_003.fcs
Id2: 12Sep2018

| Id | Alias | Total | Region | geoMeanX | StdDevX | geoMeanY | StdDevY |
|---|---|---|---|---|---|---|---|
| R2 | YG beads | 111760 | 894 | 476.99 | 143.69 | 95.07 | 33.40 |
| R3 | YG beads 520 572 | 111760 | 795 | 441.07 | 17.78 | 87.07 | 4.88 |
| R4 | YG FSC 520 | 111760 | 790 | 5306.13 | 297.05 | 440.57 | 15.03 |
| R5 | Syn | 111760 | 1050 | 42.23 | 48.02 | 47.27 | 55.02 |
| R6 | Pro | 111760 | 0 | 0.00 | 0.00 | 0.00 | 0.00 |
| R7 | Bacteria | 111760 | 30865 | 32.59 | 84.94 | 88.82 | 72.10 |
| R8 | Pro double | 111760 | 0 | 0.00 | 0.00 | 0.00 | 0.00 |

1 of 1

**Extended Data Figure 15**: Bacteria are differentiated from detritus and cyanobacteria.

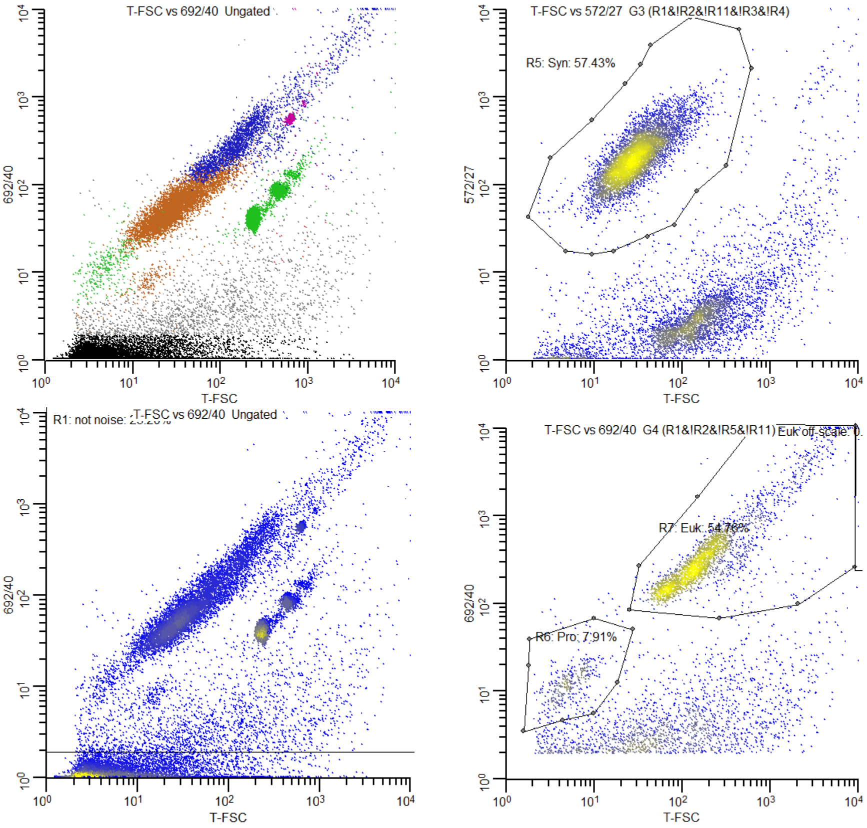


Id1: MAF_19Apr19_007.fcs
Id2: 7May2019

| Id | Alias | Total | Region | geoMeanX | StdDevX | geoMeanY | StdDevY |
|---|---|---|---|---|---|---|---|
| R2 | YG beads | 139001 | 23274 | 645.98 | 221.74 | 936.36 | 319.69 |
| R3 | YG beads 520 572 | 139001 | 21582 | 615.19 | 31.46 | 889.76 | 45.53 |
| R4 | YG FSC 692 | 139001 | 21508 | 239.58 | 13.39 | 40.53 | 3.95 |
| R5 | Syn | 139001 | 6515 | 29.82 | 29.95 | 204.92 | 175.45 |
| R6 | Pro | 139001 | 383 | 5.24 | 2.82 | 11.95 | 7.81 |
| R7 | Euk | 139001 | 2651 | 209.72 | 984.81 | 353.11 | 1518.46 |
| R10 | Rainbow T-FSC 692 | 139001 | 362 | 630.79 | 36.59 | 563.72 | 34.39 |
| R11 | Rainbow T-FSC 572 | 139001 | 401 | 660.36 | 171.32 | 89.41 | 63.73 |
| R13 | Euk off-scale | 139001 | 10 | 9922.70 | 232.98 | 6126.09 | 3725.25 |

1 of 1

**Extended Data Figure 16**: Flow cytometry gating strategy for photosynethic populations. The left column shows all data categorized (top) and as a density plot (bottom). *Synechococcus* and beads are gated first (top right) and *Prochlococcus* and picoeukaryotes are differentiated as on FSC.
